# African Swine Fever Virus CD2v protein promotes β-Interferon expression and apoptosis in swine cells

**DOI:** 10.1101/2020.11.06.372417

**Authors:** Sabal Chaulagain, Gustavo Delhon, Sushil Khatiwada, Daniel L. Rock

**Affiliations:** Department of Pathobiology, College of Veterinary Medicine, University of Illinois at Urbana-Champaign, Urbana, IL, United States of America; School of Veterinary Medicine and Biomedical Sciences, Nebraska Center for Virology, University of Nebraska-Lincoln, NE, United States of America

**Author notes:** Address correspondence to Daniel L. Rock.

**Keywords:** African swine fever virus, CD2v, Interferon-β, NF-κB, CD58, Lymphocyte, Apoptosis, Pathogenesis

## Abstract

African swine fever (ASF) is a disease of swine characterized by massive lymphocyte depletion in lymphatic tissues due to apoptosis of B and T cells, most likely triggered by proteins or factors secreted by infected adjacent macrophages. Here we describe a role for the ASF virus (ASFV) protein CD2v in apoptosis induction in lymphocytes. CD2v is a viral homolog of host CD2 that has been implicated in viral virulence and immunomodulation *in vitro*; however, its actual function remains unknown. We show that CD2v is secreted into culture medium of CD2v-expressing swine cells; and expression of-or treatment with CD2v led to significant induction of IFN-β/ISGs transcription and antiviral state. CD2v expression led to enhanced NF-κB-p65 nuclear translocation in these cultures and incubation with a NF-κB inhibitor reduced CD2v-induced NF-κB-p65 nuclear translocation and IFN-β transcription. We show that CD2v binds CD58, the natural CD2 ligand, and that CD58 siRNA knock-down results in significant reduction in NF-κB-p65 nuclear translocation and IFN-β transcription. Treatment of swine PBMC with purified CD2v led to enhanced NF-κB-p65 nuclear translocation and induction of IFN-β transcription. Further, induction of caspase-3 and PARP1 cleavage was observed in these swine PBMC at later times, providing a mechanism for CD2v-induced apoptosis of lymphocytes. Finally, IFN-β induction and NF-κB activation was inhibited in swine PBMC treated with purified CD2v pre-incubated with antibodies against CD2v. Overall, our results indicate that ASFV CD2v is an immunomodulatory protein that, by promoting lymphocyte apoptosis, may contribute to bystander lymphocyte depletion observed during ASFV infection in pigs.

**IMPORTANCE:** ASF, a severe hemorrhagic disease of domestic swine, represents a significant economic threat to swine industry worldwide. One critical pathological event observed in pigs infected with virulent isolates is an extensive destruction of lymphoid tissue and massive lymphocyte depletion. However, viral factor/s involved in this event are yet to be identified. Here we show that, by inducing NF-κB-dependent IFN signaling, ASFV CD2v is able to promote apoptosis in swine PBMC. We propose that CD2v released by ASFV-infected macrophages contributes to the massive depletion of lymphocytes observed in lymphoid tissues of infected pigs. Results here improve our understanding of ASFV pathogenesis and will encourage novel intervention approaches.

## INTRODUCTION

African swine fever (ASF) is a severe hemorrhagic disease of swine caused by ASF virus (ASFV), a complex enveloped DNA virus which is currently the sole member of *Asfarviridae* family and the only known DNA arbovirus. The disease is endemic to sub-Saharan African countries where virus cycles between bushpigs and warthogs, and *Ornithodorus sp.* ticks (1–4). While infection of wild pigs is asymptomatic, infection of domestic pigs can result in mortality approaching 100%. Clinical forms of ASF disease range from per-acute to chronic depending on the virus strain and host factors (5–7). Lesions are most notably seen in lymphoid organs, and are characterized by massive lymphocyte depletion and lymphatic tissue destruction (6–10). Although ASF virus replicates in macrophages (11–14), infection with virulent ASFV causes marked apoptosis in B and T cells, which are not targets for viral replication (3, 15–19). It has been suggested that lymphocyte apoptosis is most likely induced by proteins or factors secreted or released by infected macrophages (19–22). Currently, no ASFV protein function has been associated with apoptosis induction in lymphocytes.

Interferons (IFNs) provide the first line of defense against virus infection. There are three different types of IFNs, types I, II and III. Type I IFNs consist of 13 IFN-α subtypes and IFN-β. Type II and III IFN consists of IFN-γ and 3 subtypes of IFN-λ, respectively. Nearly all cell types produce type I IFNs when host pathogen recognition receptors (PRRs) bind different pathogen associated molecular patterns (PAMPs) (23). IFNs exert their effects by signaling through their specific receptors, leading to transcriptional induction of interferon stimulated genes (ISGs). ISGs mediate the anti-viral, anti-proliferative and immunomodulatory functions of IFN (24). Uncontrolled high levels of type I IFN during viral infection might be detrimental to the host as it has been associated with immunopathology, immunosuppression, enhanced pathology and disease progression (25). The importance of interferon for control of virus infections is reflected by the myriad of genes encoded by different types of viruses that target interferon responses (26, 27).

IFN-β induction is mediated by two major groups of transcription factors, nuclear factor-kB (NF-κB) and IFN-regulatory factors (IRFs) (28–32). Canonical NF-κB activation involves inhibitor kappa-β kinase (IKK) complex activation that leads to phosphorylation and degradation of inhibitor kappa-β (IκB) proteins resulting in NF-κB nuclear translocation and subsequent transcriptional induction of pro-inflammatory genes, including IFN-β (33–38). Lack of NF-κB activity leads to decreased expression of IFN-β in cells (39, 40). Because of the central role of NF-κB in various antiviral responses, viruses target multiple steps on NF-κB activation, from PRR recognition to NF-κB mediated gene transcription (41).

ASFV genome encodes several genes that function to interfere with NF-κB and IFN pathways in macrophages. For example, genes of the multi-gene family 360 and 530 (MGF360/MGF530) suppress IFN induction through yet unidentified mechanisms; DP96 inhibits the cGAS-STING-TBK pathway initiated by viral DNA, leading to inhibition of NF-κB /IRF-3-mediated IFN-β induction; A238L inhibits NF-κB activation by binding to the NF-κB subunit RelA, and by inhibiting the acetylation of NF-κB-p65 by p300;and I329L inhibits dsRNA-induced NF-κB and IRF3 activation (42–45). Although virulent ASFV induces very low level of IFNs inconsistently during infection of macrophages *in vitro* (46–51), pigs infected with virulent ASFV show high levels of IFNs in serum and enhanced levels of cytokines (TNFα, IL-1α, IL-1β and IL-6) in serum and organs (19–22, 52, 53). This suggests a role/s of IFNs of ASFV pathogenesis in pigs.

Uncharacterized soluble factors released by ASFV-infected macrophages have been shown to inhibit proliferation of swine lymphocytes in response to lectins (54). CD2v, the ASFV hemagglutinin and homolog of host T cell and NK cell surface antigen CD2, has been shown to inhibit the proliferation of lymphocytes in response to lectins, suggesting that CD2v has immunosuppressive activity *in vitro* (55). CD2v contains all the domains present in cellular CD2 and some of the residues involved in binding to its natural ligand, CD58 (56, 57). CD2-CD58 interaction has been shown to activate cellular kinases in lymphocytes; however, the involvement of NF-κB signaling downstream CD2-CD58 interaction has not been shown (58–67).

The aim of this study is to investigate immunomodulatory function/s of ASFV CD2v and define a role of CD2v in induction of lymphocyte apoptosis. We show here that expression of-or treatment with-CD2v leads to induction of IFN-β, ISGs and the antiviral state in swine cells. We also show that CD2v-induced IFN-β expression requires NF-κB activation and CD2v-CD58 interaction. Importantly, CD2v treatment induces apoptosis in swine PBMC.

## RESULTS

### ASFV CD2v localizes in perinuclear region, cytoplasm and cell membrane of PK15 cells and is secreted into the culture supernatant

CD2v is an ASFV structural transmembrane glycoprotein homologous to CD2, a cell adhesion molecule expressed by T- and NK-cells (56, 57). CD2v encodes for a protein of 370 amino acids, with a predicted molecular weight of 42 kDa and is expressed on the surface of ASFV-infected macrophages. CD2v mediates hemadsorption of swine red blood cells (RBCs) (56, 57) and, together with viral C-type lectin, has a role in hemadsorption inhibition (HAI) serotype specificity (71).

To examine the subcellular localization and expression kinetics of CD2v, PK15 cells were mock transfected or transfected with pCMV plasmid expressing C-terminally HA-tagged CD2v (CD2v-HA) and examined at various times post-transfection (pt) by confocal microscopy. CD2v was observed adjacent to the nucleus at 2 h pt, and in the cell membrane, perinuclear area and cytoplasmic vesicles at later times pt (Fig. 1A). To confirm that CD2v expressed by PK15 cells is capable of hemadsorption, PK15 cells were transfected with pEmpty-HA (control plasmid) or pCD2v-HA and incubated with swine red blood cells (RBCs) 24 h later. PK15 transfected with pCD2v-HA but not with control plasmid hemadsorbed swine RBCs as evidenced by rosette formation (Fig. 1B, arrowheads).

**FIG 1.**
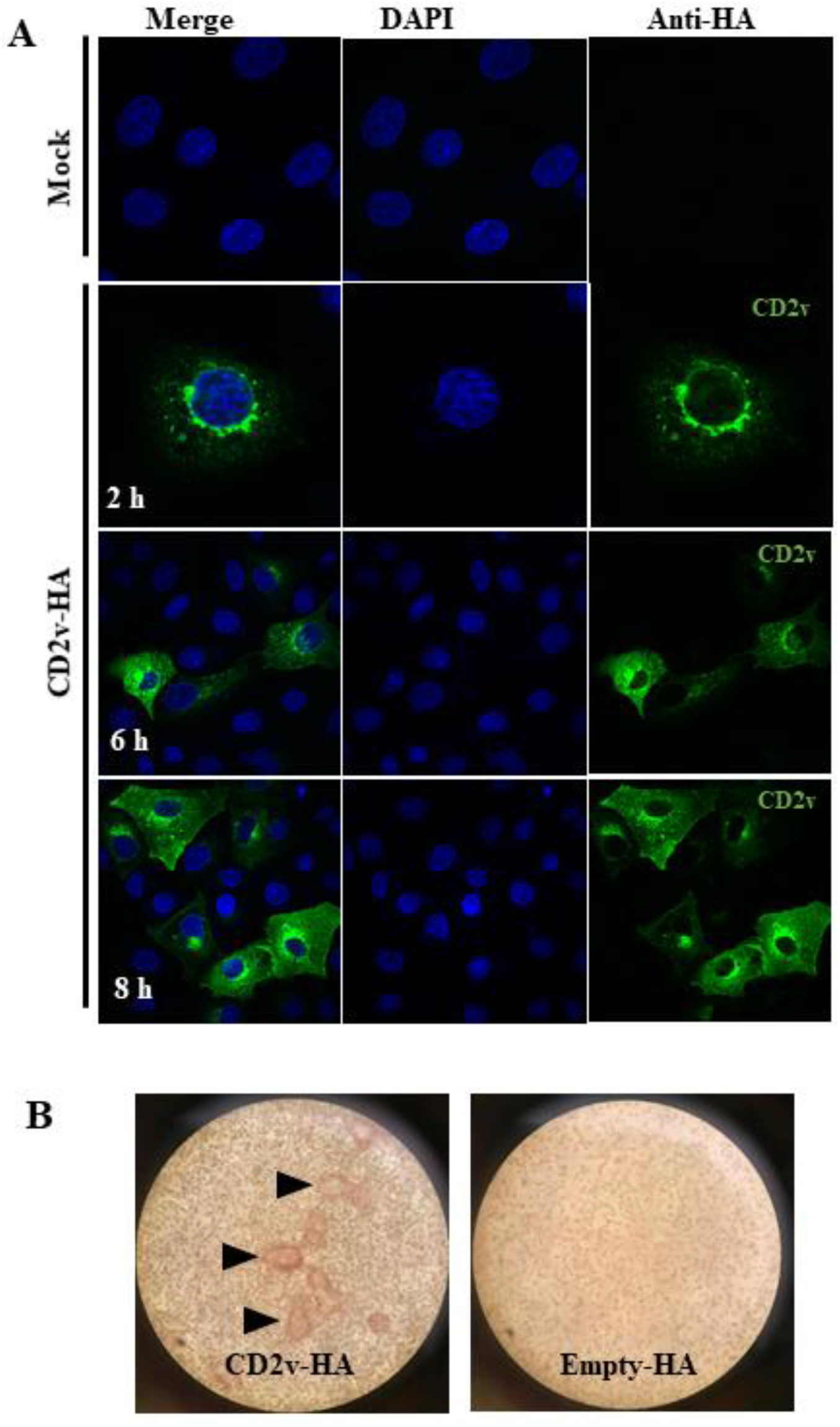
Subcellular localization and characterization of ASFV CD2v protein. (A) PK15 cells were mock transfected or transfected with a plasmid expressing HA-tagged ASFV CD2v, fixed at various times post transfection (pt), incubated with anti-HA primary antibody, sequentially stained with Alexa fluor 488-labeled secondary antibody and DAPI, and examined by confocal microscopy. Results are representative of two independent experiments. (B) CD2v-HA-expressing PK15 cells hemadsorbed swine red blood cells (RBCs). PK15 cells were transfected with plasmid pCD2v-HA for 24 h, incubated with swine RBCs overnight, and observed with the microscope (X100). CD2v-dependent rosetting is indicated by the arrowheads. Results are representative of two independent experiments.

The expression kinetics of CD2v was assessed by Western blot after transfection of PK15 cells with CD2v-HA. Two major protein species of approximately 100 kDa and 25 kDa, and a less-abundant 15 kDa species were detected at 6 h pt, with increasing protein levels observed at later time points (Fig. 2A). A similar expression pattern was observed in 293T cells and Vero cells (Fig. S1 in supplemental material). The observed molecular weight of the full length protein was approximately 58 kDa higher than predicted. A single 42 kDa band was detected when CD2v was expressed in presence of tunicamycin, an inhibitor of N-linked glycosylation, confirming that the protein is heavily modified through N-linked glycosylation (Fig. S2 in supplemental material) (72). Because the 25 kDa species is absent in presence of tunicamycin, we speculate that the 25 kDa protein product might result from processing of the full length protein in the endoplasmic reticulum. A study by Goatley and Dixon (2011) has shown that CD2v in virus-infected cells is cleaved in the endoplasmic reticulum or Golgi compartments. A faint 100 kDa and a predominant 25 kDa CD2v band were detected in the culture supernatant of PK15 cells 24 h post transfection with pCD2v-HA, indicating that CD2v is secreted into the culture supernatant (Fig. 2B).

**FIG 2.**
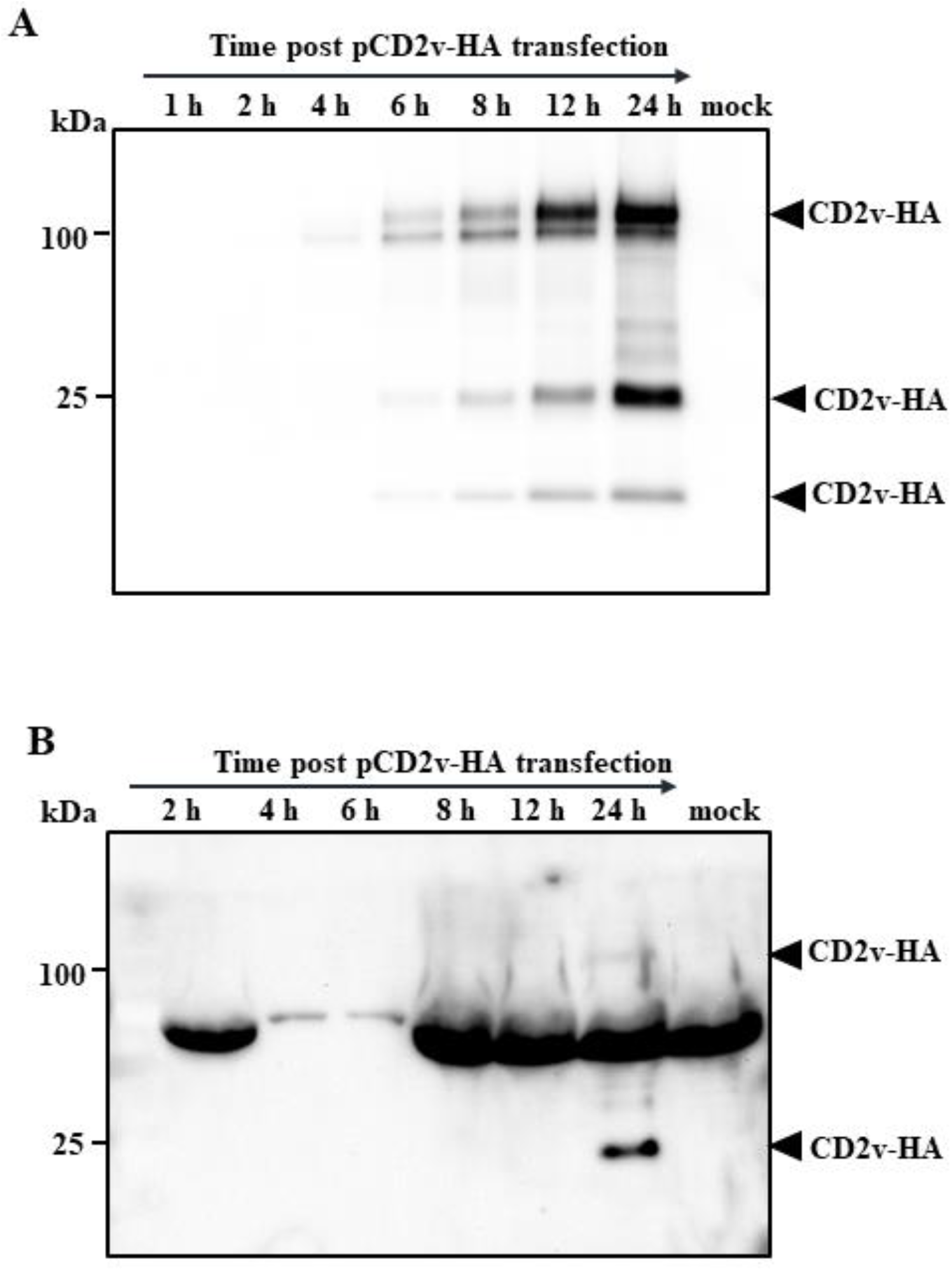
Expression kinetics and secretion of CD2v. (A) PK15 cells mock transfected or transfected with plasmid pCD2v-HA were harvested at the indicated times post transfection. Total cell protein extracts were resolved by SDS-PAGE, blotted and incubated with antibodies against HA. Results are representative of two independent experiments. (B) Detection of CD2v in culture supernatant of PK15 cells transiently expressing CD2v. PK15 cells were transfected as above, and supernatants harvested at various times pt. Cleared supernatants (50 μl) were resolved by SDS-PAGE and analyzed by Western blot using antibodies against HA. Results are representative of two independent experiments.

Overall, our results are in agreement with previous studies on CD2v expression in ASFV-infected cells (56, 72, 73). In addition, this is the first study showing that CD2v is secreted into the culture medium, as previously speculated by others (72, 74).

### Expression of ASFV CD2v in PK15 cells induces IFN-β and ISGs transcription, and the antiviral state

Preliminary RNA-Seq experiments were conducted to examine the effect of secreted CD2v on cellular gene transcription. PK15 cells were incubated with CD2v-containing supernatant (1:2 dilution) or supernatant from cells transfected with pEmpty-HA (control plasmid), and total RNA was collected at 1 h, 2 h and 3 h post-treatment. RNA-Seq analysis showed upregulation of several interferon-stimulated genes (ISGs), including *MX1, OAS1,* and *IRF9* at 2 h post treatment with further increase at 3 h, suggesting a potential role of CD2v in IFN-β signaling (data not shown). To assess the effect of CD2v in IFN-β and ISG transcription, PK15 cells were transfected with pCD2v-HA or control plasmids pEmpty-HA and pORFV120-Flag, the latter encoding for Orf virus protein ORFV120 and IFN-β transcription was assessed by RT-PCR. Compared to controls, cells transfected with pCD2v-HA plasmid showed significant upregulation of IFN-β (5.6-fold) as early as 6 h pt and similar upregulation was observed at all other subsequent time points (Fig. 3A). Consistent with upregulation of IFN-β, significant increase of ISGs *MX1* (17.7-fold) and *OAS1* (12.8-fold) transcription was observed at 30 h pt with pCD2v-HA plasmid compared to controls (Fig. 3B and C).

**FIG 3.**
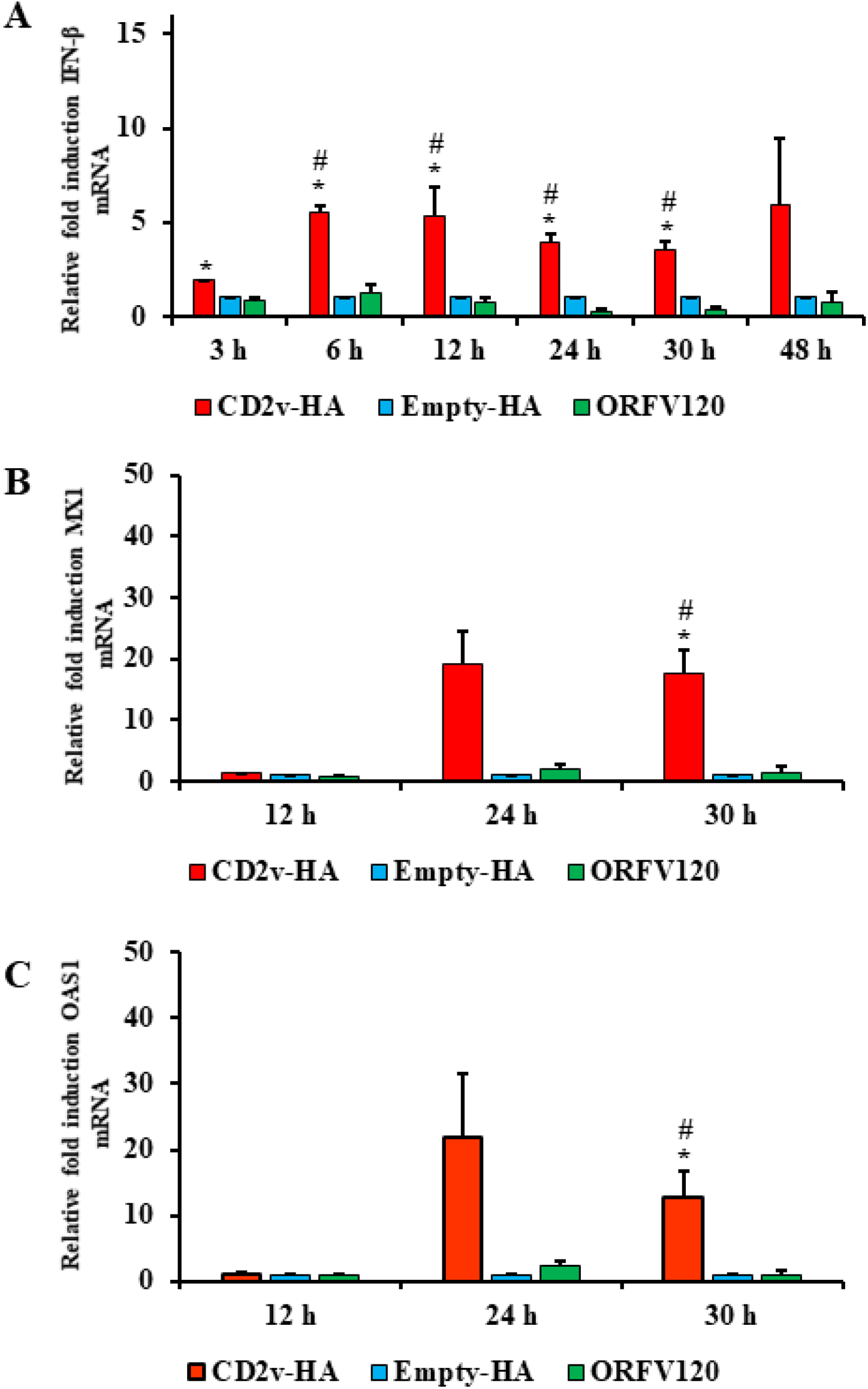
Induction of IFN-β and ISGs in CD2v-expressing cells. PK15 cells were transfected with plasmids pCD2v-HA, pEmpty-HA or pORFV120-Flag, and transcription of IFN-β and selected ISGs were assessed by real-time PCR. (A) Fold changes of IFN-β levels in cells transfected with pCD2v-HA relative to pEmpty-HA and pORFV120-Flag at various times post transfection (pt). Results are average mRNA levels from three independent experiments. *P*-values relative to Empty-HA and ORFV120 were 0.007 and 0.1 (3 h); 0.04 and 0.01 (6 h); 0.016 and 0.017 (12 h); 0.015 and 0.006 (24 h); and 0.019 and 0.006 (30 h), respectively. (B and C) Fold changes of ISG mRNA levels at 12 h, 24 h and 30 h pt. Results are average mRNA levels from three independent experiments. *P*-values relative to Empty-HA and ORFV120 at 30 h pt were 0.02 and 0.018 for *MX1* (B) and 0.04 and 0.037 for OAS1 (C). * and # denote statistical significance compared to Empty-HA and ORFV120, respectively.

To investigate the functional significance of IFN-β and ISGs induction by CD2v, the antiviral state of cells was examined using an IFN bioassay. PK15 cells were transfected with pCD2v-HA or control plasmids (Empty-HA vector or plasmids expressing Orf virus proteins ORFV120 and ORFV113), and then infected with reporter VSV^GFP^ (50 PFU/well) at various times pt. PK15 cells transfected with pCD2v-HA but not with control plasmids showed inhibition of VSV^GFP^ replication as determined by both fluorescent microscopy and flow cytometry at 12 h, 24 h and 30 h pt (Fig. 4A and B). Inhibition of VSV^GFP^ replication was also observed after treatment of PK15 cells with a two-fold dilution of culture supernatants from cells transfected with Poly (I:C) (up to dilution 1:8) and pCD2v-HA (up to dilution 1:4) but not with control plasmids (Fig. 4C). These data indicate that expression of-or treatment with-CD2v in PK15 cells leads to induction of IFN-β, ISGs and an antiviral state.

**FIG 4.**
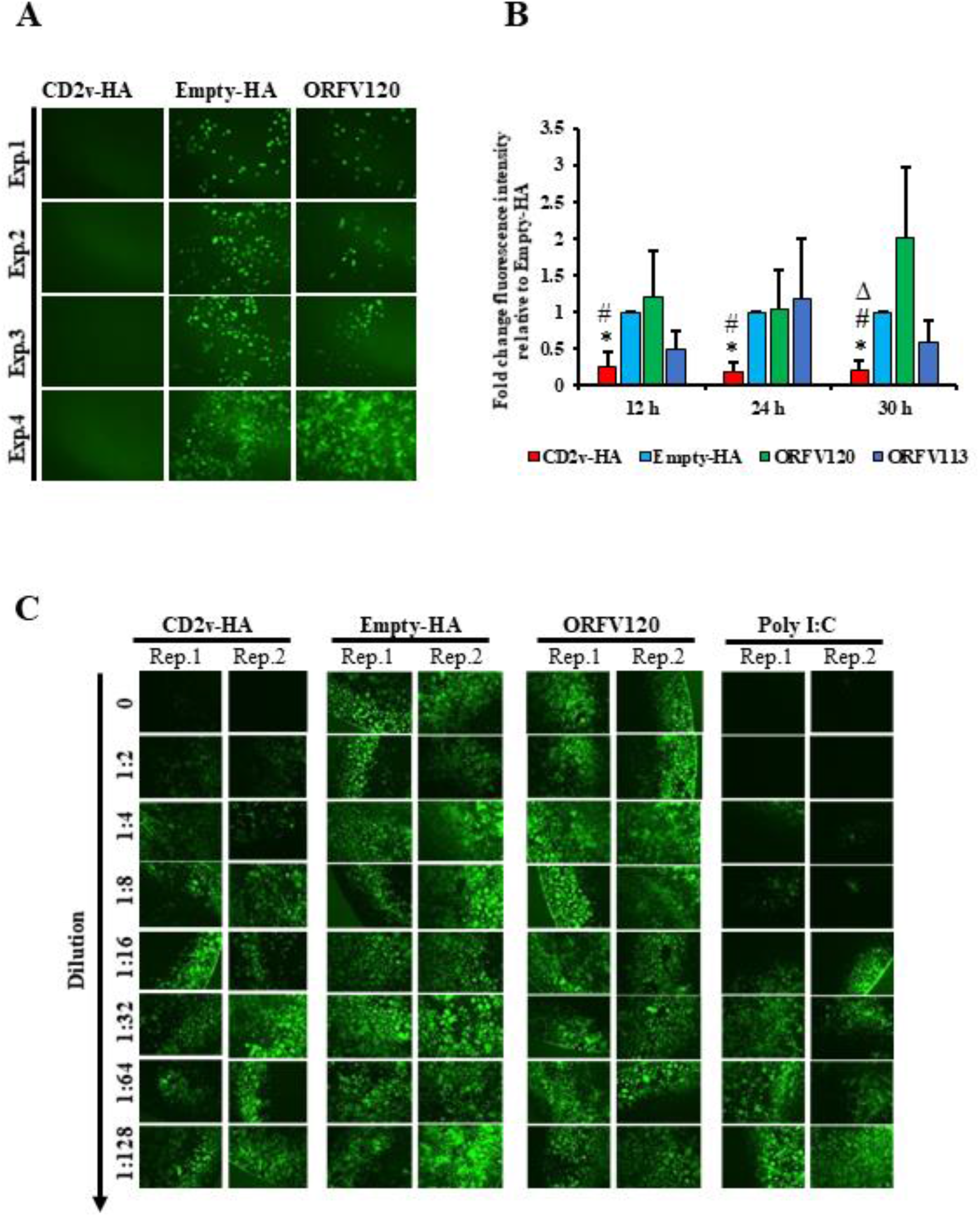
Induction of antiviral state in CD2v-expressing cells. PK15 cells were transfected with pCD2v-HA, pEmpty-HA, pORFV120-Flag or pORFV113-Flag. At 12 h, 24 h or 30 h post transfection, cultures were infected with vesicular stomatitis virus expressing GFP (VSV^GFP^, 50 PFU/well). (A) Fluorescence microscopy images taken at 16 h post infection. Note decreased VSV replication in cells transfected with pCD2v-HA relative to controls. Results representative of four independent experiments. Exp., denotes experimental replicates. (B) Mean GFP fluorescence measured by flow cytometry at 16 h post infection. Results are mean values of four independent experiments. *P*-values relative to transfection with plasmids pEmpty-HA, pORFV120-Flag and pORFV113-Flag were 0.0064, 0.042 and 0.096 (12 h); 0.0002, 0.041 and 0.07 (24 h); and 0.0002, 0.02 and 0.02 (30 h), respectively. *, # and ∆ denote statistical significance compared to Empty-HA, ORFV120 and ORFV113, respectively. (C) PK15 cells were treated with supernatants obtained from cultures transfected with pCD2v-HA, pEmpty-HA or pORFV120-Flag, or with poly I: C, and infected with VSV^GFP^ (50 PFU/well) 30 h post treatment. Fluorescence microscopy images taken at 16 h post infection are shown. Results representative of two independent experiments. Rep., denotes technical replicates.

### Induction of IFN-β by ASFV CD2v depends on NF-κB activation

NF-κB and IRF3 are two important transcription factors involved in IFN-β induction (28–32) that translocate to the nucleus upon activation. To assess whether CD2v expression affects NF-κB-p65 and IRF3 nuclear translocation, PK15 cells were transfected with pCD2v-HA or control plasmids pORFV120-Flag and pORFV113-Flag, and examined by IFA at various times pt. Enhanced NF-κB-p65 nuclear translocation was observed in CD2v expressing cells at all times pt compared to controls (Fig. 5A and B). In contrast, IRF3 nuclear translocation was not observed (data not shown). To confirm activation of the NF-κB pathway, PK15 cultures were transfected with pCD2v-HA or pORFV113-Flag (control), and the mean fluorescence intensity (MFI) of phosphorylated NF-κB (pNF-κB S536) was examined at 3 h pt by flow cytometry. Consistent with the nuclear translocation results, significantly increased MFI values (1.8-fold) were observed in CD2v-expressing cells compared to the control (Fig. 5C).

**FIG 5.**
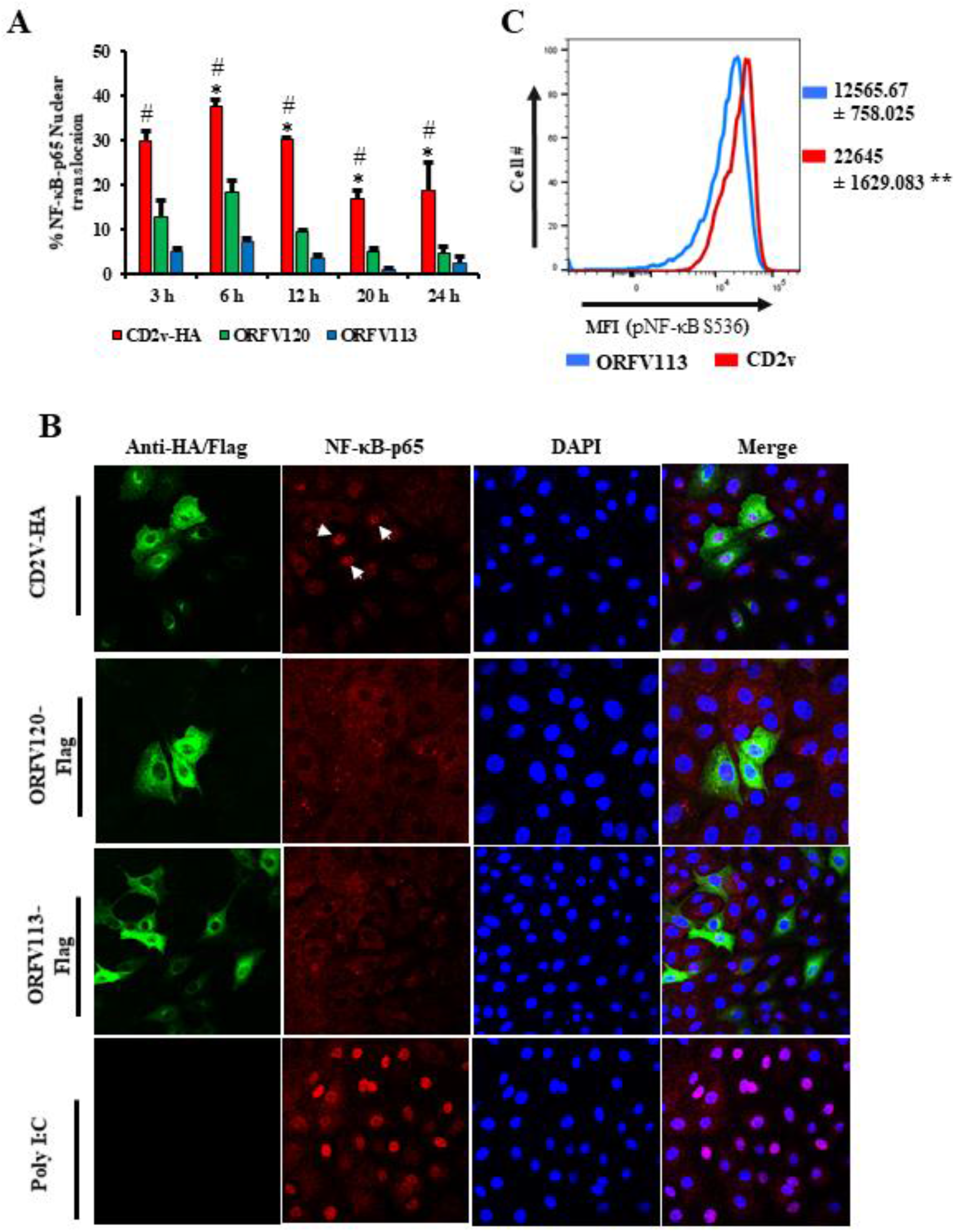
Induction of NF-κB-p65 nuclear translocation in CD2v-expressing cells. PK15 cells were transfected with pCD2v-HA, pORFV120-Flag or pORFV113-Flag, processed for immunofluorescence using primary antibodies against HA or Flag and NF-κB-p65, and secondary antibodies Alexa fluor 488 to detect CD2v, ORFV120 or ORFV113 and Alexa fluor 594 to detect NF-κB-p65, and counterstained with DAPI. Cells were counted from 15 random fields/slide (approximately 100 cells/slide) and results, shown as percentage of cells with nuclear NF-κB-p65, are mean values from three independent experiments. (A) Percentage of NF-κB-p65 nuclear translocation following the different treatments. *P*-values relative to ORFV120 and ORFV113 were 0.056 and 0.009 (3 h); 0.01 and 0.0006 (6 h); 0.019 and 0.02 (12 h); 0.005 and 0.004 (20 h); and 0.016 and 0.009 (24 h), respectively. * and # denote statistical significance compared to ORFV120 and ORFV113, respectively. (B) Confocal microscopy images showing NF-κB-p65 nuclear translocation at 3 h post transfection (pt; arrows). Green, CD2v or ORFV120 or ORFV113; Red, NF-κB-p65; Blue, DAPI. (C) Mean fluorescence intensity (MFI) of phosphorylated NF-κB (S536) fluorescence measured by flow cytometry in CD2v and ORFV113 expressing cells at 3 h pt. Results representative of three independent experiments. *P*-value relative to ORFV113 is 0.0082.

To assess whether inhibition of NF-κB affects IFN-β induction by CD2v, PK15 cells were pretreated with the NF-κB inhibitor parthenolide (1μM) or vehicle control (DMSO) for one hour, transfected with pCD2v-HA in presence of the inhibitor (1μM) or vehicle, and assessed for NF-κB-p65 nuclear translocation at 3 h pt by confocal microscopy. Percentage of NF-κB-p65 nuclear translocation in CD2v-expressing cells in presence of parthenolide was significantly reduced (49.3%) compared to DMSO (Fig. 6A and B).

**FIG 6.**
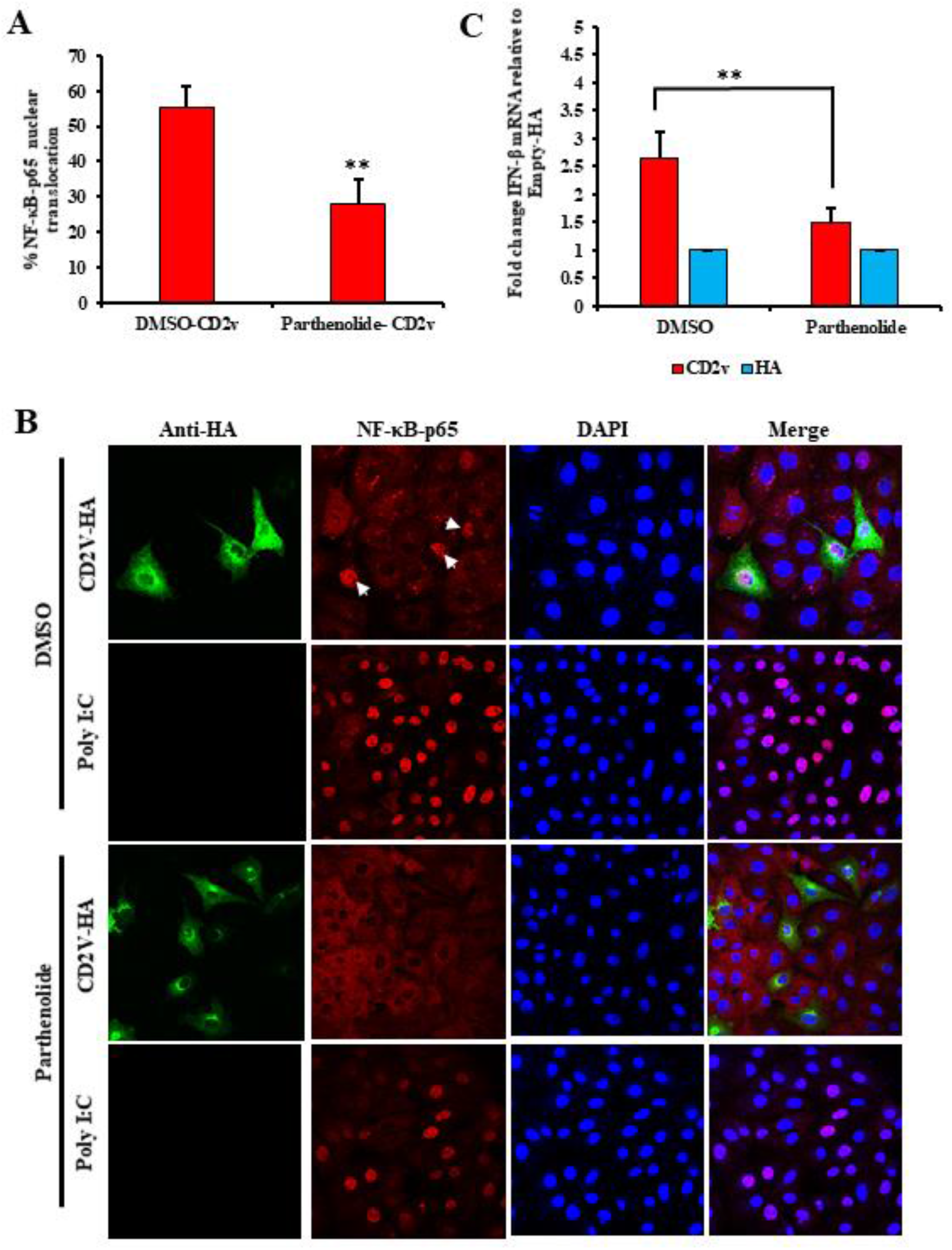
CD2v expression induces IFN-β transcription in NF-κB dependent manner in PK15 cells. PK15 cells pretreated with the NF-κB inhibitor parthenolide (1 μM) or DMSO (vehicle control) for one hour, were transfected with pCD2v-HA, fixed at 3 h post transfection and processed for immunofluorescence with antibodies against HA and NF-κB-p65. Cells were counted from 15 random fields/slide (approximately 100 cells/slide) (A) Percentage of CD2v-expressing cells with nuclear NF-κB-p65. Results are expressed as mean values from three independent experiments (*P* = 0.001). (B) Representative confocal images of cells treated as above. Green, CD2v; Red, NF-κB-p65; Blue, DAPI. Arrows indicate nuclear NF-κB-p65. (C) PK15 cells treated with parthenolide (1 μM) or DMSO for one hour, were transfected with pCD2v-HA in presence or absence of parthenolide. Total RNA was extracted at 6 h pt, cDNA prepared and IFN-β transcription assessed by RT-PCR. Fold changes relative to Empty-HA and data are means from four independent experiments (*P* = 0.008). (*, *P*<0.05; **, *P*<0.01).

To investigate the effect of NF-κB inhibition on CD2v-mediated IFN-β induction, PK15 cells were pretreated for one hour with parthenolide (1μM) or DMSO (vehicle control), transfected with pCD2v-HA or pEmpty-HA (control) for 6 h in presence of parthenolide (1μM) or DMSO, and assessed for IFN-β transcription by RT-PCR. Significant reduction in IFN-β transcription was observed in cells transiently expressing CD2v in presence of parthenolide (1.5-fold) compared to cells expressing CD2v in presence of vehicle alone (2.6-fold) (Fig. 6C). Together, results above indicate that induction of IFN-β by ASFV CD2v is mediated by NF-κB activation.

### NF-κB activation and IFN-β induction by ASFV CD2v is mediated by interaction of CD2v with host CD58

CD2v contains all the domains present in cellular CD2 and some of the residues involved in binding to the CD58, the natural CD2 ligand (56, 57). To study the potential interaction between CD2v and CD58, PK15 cells were co-transfected with pEmpty-HA vector or pCD2v-HA and porcine pCD58-Flag, and cell lysates were prepared at 8 h pt for reciprocal co-immunoprecipitation with anti-Flag or anti-HA antibodies as described in materials and methods. Figure 7A shows that CD2v and porcine CD58 reciprocally co-immunoprecipitate.

**FIG 7.**
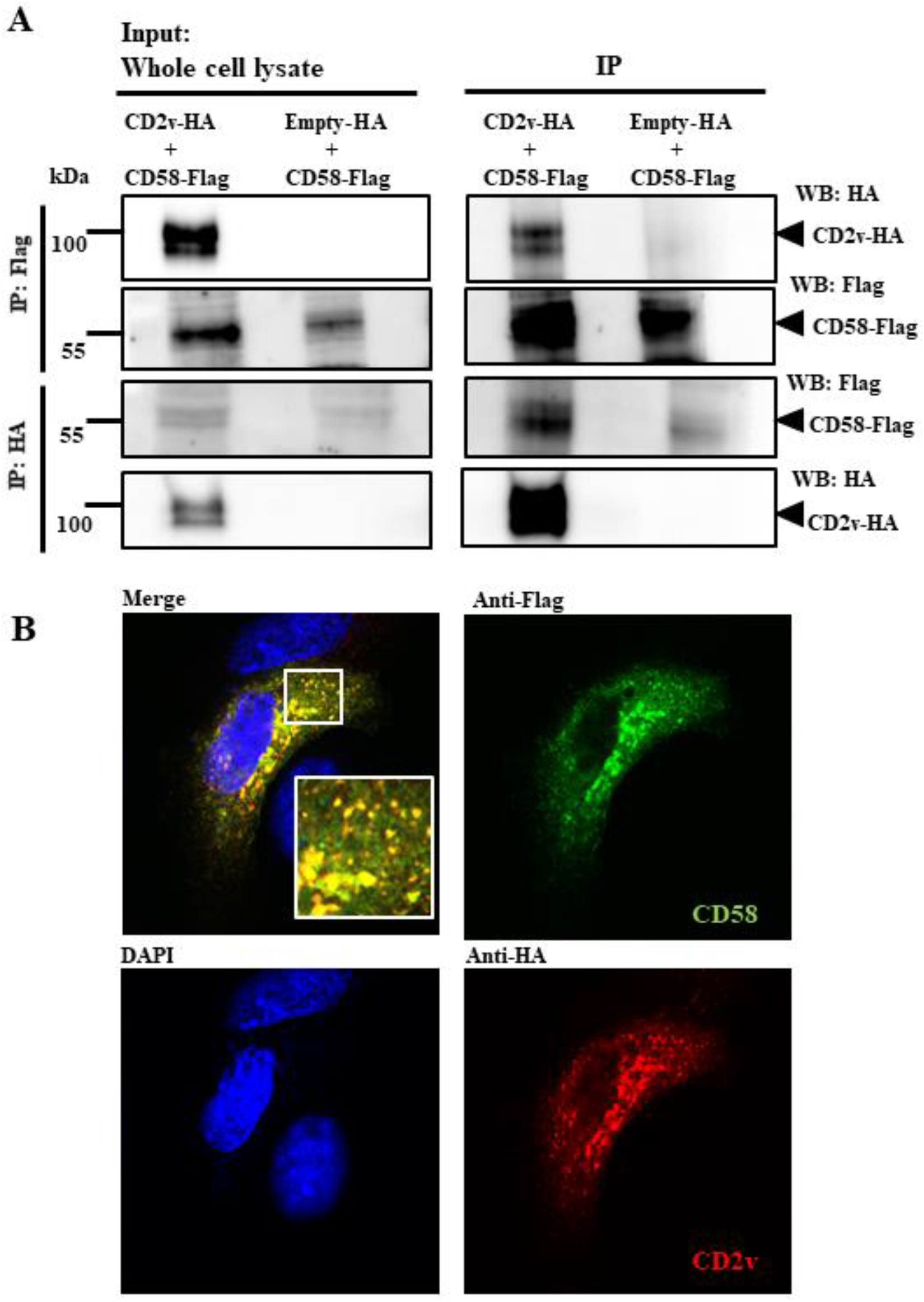
Interaction between CD2v and porcine CD58. (A) For co-immunoprecipitation experiments, PK15 cells were co-transfected with plasmids pCD58-Flag and pCD2v-HA, or pCD58-Flag and pEmpty-HA (control) and harvested at 8 h post transfection (pt). Whole cell lysates (left) and extracts immunoprecipitated with anti-Flag antibodies (IP, two top panels) or anti-HA (IP, two bottom panels) were examined by Western blotting with antibodies directed against proteins indicated on the right. Results are representative of three independent experiments. (B) For co-localization studies, PK15 cells were co-transfected with pCD58-Flag and pCD2v-HA, fixed at 24 h pt, incubated with mouse anti-Flag and rabbit anti-HA primary antibodies, washed, and incubated with secondary antibodies (Alexa-fluor 488-labeled anti-mouse and Alexa-fluor 594-labeled anti-rabbit). Cells were counterstained with DAPI, and examined with the confocal microscope. Results are representative of three independent experiments. Insets show magnified areas of the field.

To confirm CD2v-CD58 interaction, PK15 cells were co-transfected with pCD2v-HA and pCD58-Flag, and co-localization of proteins was examined using confocal microscopy. Strong overlap of signals was observed (Fig. 7B). Similarly, strong reciprocal co-immunoprecipitation and co-localization of CD2v with endogenous human CD58 was observed in 293T cells transfected with pCD2v-HA, using anti-HA or anti-hu-CD58 antibodies (Fig. 8A and B). Given that: 1) CD2v interacts with CD58 (Fig. 7 and Fig. 8) and 2) host CD2-CD58 interaction activates downstream cellular kinases (58–67), we examined the involvement of CD2v-CD58 interaction in CD2v-mediated NF-κB activation and IFN-β activation.

**FIG 8.**
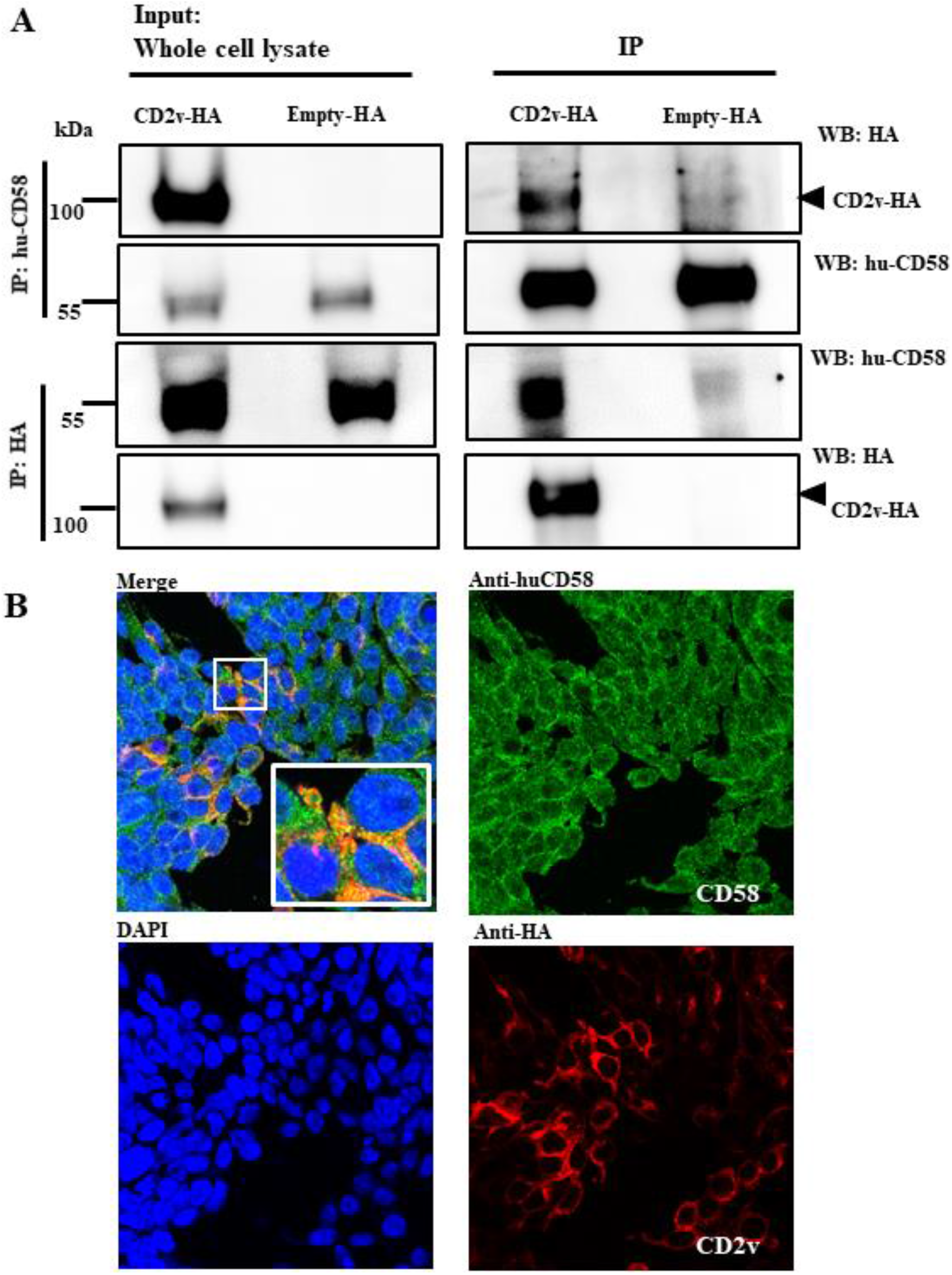
Interaction between CD2v and human CD58. (A) For co-immunoprecipitation, 293T cells were transfected with pCD2v-HA or pEmpty-HA (control) and harvested at 8 h pt. Whole cell lysates (left) and extracts immunoprecipitated with mouse anti-huCD58 antibodies (IP, two top panels) or anti-HA (IP, two bottom panels) were examined by Western blotting with antibodies directed against proteins indicated on the right. Results are representative of three independent experiments. (B) For co-localization, 293T cells were transfected with pCD2v-HA, fixed at 24 h pt, incubated with mouse anti-huCD58 and rabbit anti-HA primary antibodies, washed, and incubated with Alexa-fluor 488-labeled anti-mouse and Alexa-fluor 594-labeled anti-rabbit secondary antibodies. Cells were counter stained with DAPI, and examined with the confocal microscope. Results are representative of three independent experiments. Insets show magnified areas of the field.

To evaluate the effect of CD58 downregulation on CD2v-mediated NF-κB activation, PK15 cells were transfected with siRNAs targeting porcine CD58 or control siRNA as described in materials and methods. CD58 transcript knock-down of approximately 55% was routinely obtained when PK15 cells were transfected with CD58 siRNA compared to the negative control (Fig. 9A). Twenty-four hours after siRNA treatment, cells were transfected with pCD2v-HA for 3 h, and NF-κB-p65 nuclear translocation was assessed by confocal microscopy. Significant reduction in NF-κB-p65 nuclear translocation (60%) in CD2v expressing cells was observed in PK15 cells with reduced CD58 transcript compared to the control (Fig. 9B, C and D).

**FIG 9.**
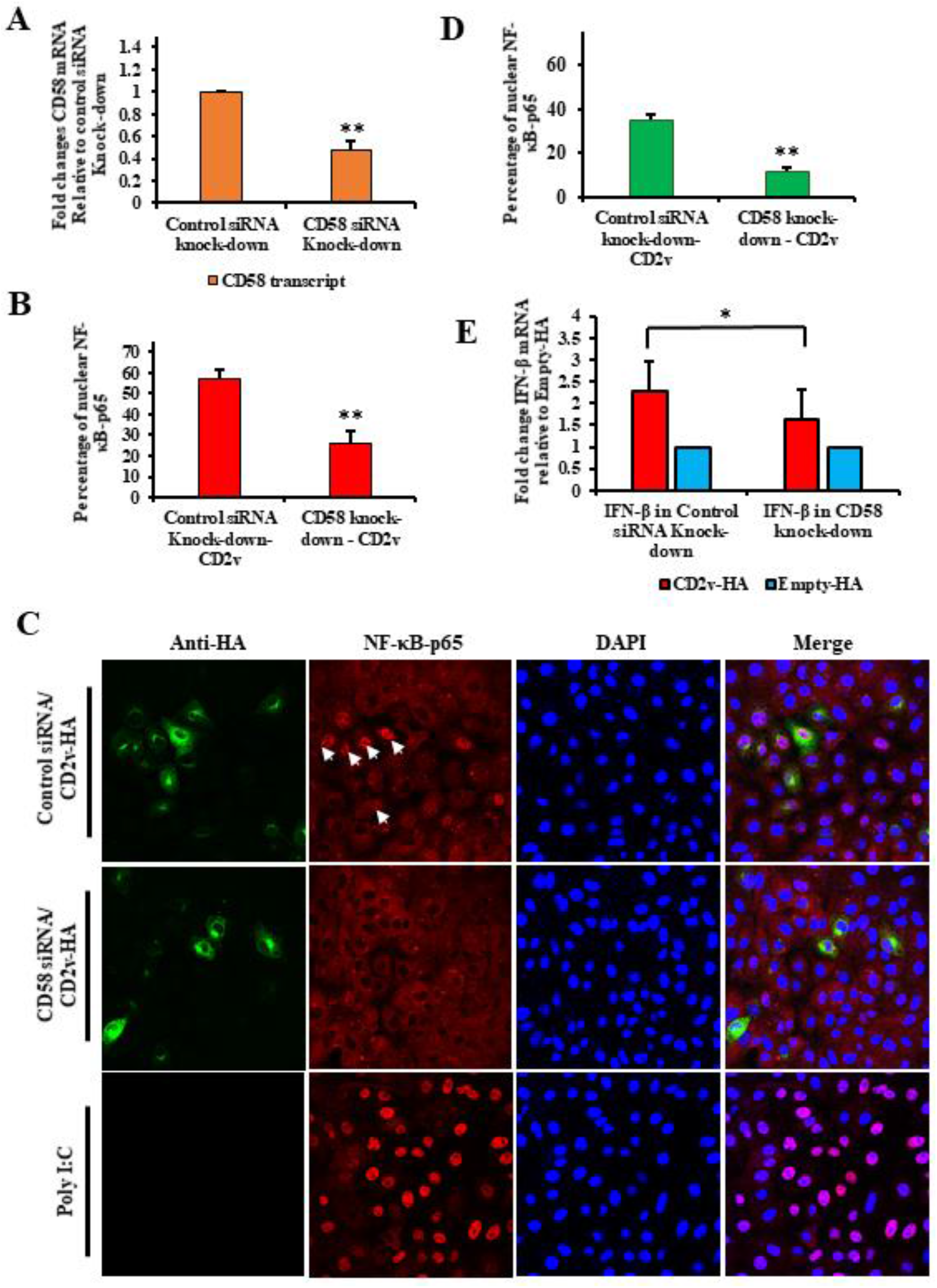
CD2v-CD58 interaction affects CD2v-mediated NF-kB-p65 nuclear translocation and IFN-β induction. (A) siRNA knock down of CD58. PK15 cells were transfected with CD58 siRNA or siRNA universal negative control, total RNA was extracted at 24 h pt, cDNA prepared and CD58 transcription assessed by RT-PCR. Results are mean of five independent experiments (*P* = 0.0002). (B) NF-κB-p65 nuclear translocation. PK15 cells were sequentially transfected with CD58 siRNA or siRNA universal negative control and pCD2v-HA or pEmpty-HA, fixed at 3 h pt, and processed for detection of NF-κB-p65 by immunofluorescence. Cells were counted from 15 random fields/slide (approximately 100 cells/slide) and results are shown as percentage of CD2v-expressing cells with nuclear NF-κB-p65. Results are mean of three independent experiments (*P* = 0.0035). (C) Percentage of CD2v-expressing cells with strong nuclear NF-κB-p65 translocation (*P* = 0.0015). (D) Representative images of NF-κB-p65 nuclear translocation under conditions outlined in A. Green, CD2v; Red, NF-κB-p65; Blue, DAPI. Arrows indicate nuclear NF-κB-p65. (E) IFN-β induction. PK15 cells were sequentially transfected with CD58 siRNA or siRNA universal negative control, and pCD2v-HA or pEmpty-HA. Total RNA was extracted at 6 h pt, cDNA prepared and transcription of IFN-β assessed by RT-PCR. Fold changes are relative to Empty-HA and data are mean mRNA levels from eight independent experiments (*P =* 0.025). (*, *P*<0.05 and **, *P*<0.01).

To investigate involvement of CD58-CD2v interaction in CD2v-mediated IFN-β induction, siRNA knock down experiments and CD2v-HA/control transfections were performed as described above. Significant reduction of IFN-β transcription was observed 6 h pt with CD2v-HA in PK15 cells with reduced CD58 transcript levels (1.6-fold) compared to the control (2.3-fold) (Fig. 9E).

### Purified CD2v induces NF-κB-p65 nuclear translocation and IFN-β transcription in swine PBMC cultures

Uncharacterized soluble factors released by ASFV-infected macrophages have been shown to inhibit proliferation of swine lymphocytes in response to lectins (54) and CD2v has been shown to be involved in inhibition of mitogen-induced proliferation of bystander lymphocytes in virus-infected swine PBMC cultures (55). This and our observations that CD2v-expressing PK15 cells secrete CD2v into the culture supernatant, and that cells treated with-or expressing CD2v upregulate IFN-β and ISGs led us to hypothesize that 1-secreted CD2v induces IFN-β and ISGs expression in lymphocytes and 2-induction involves NF-κB-p65 nuclear translocation.

To examine NF-κB-p65 nuclear translocation in swine lymphocytes, swine PBMC were treated with purified CD2v or purified control for 1.5 or 2 h, processed for IFA and deposited onto glass slides with a cytospin. Confocal microscopy analysis showed enhanced NF-κB-p65 nuclear translocation in swine PBMC treated with purified CD2v (2.8-fold at 1.5 h; 1.6-fold at 2 h) compared to control treatment (Fig. 10A and B).

**FIG 10.**
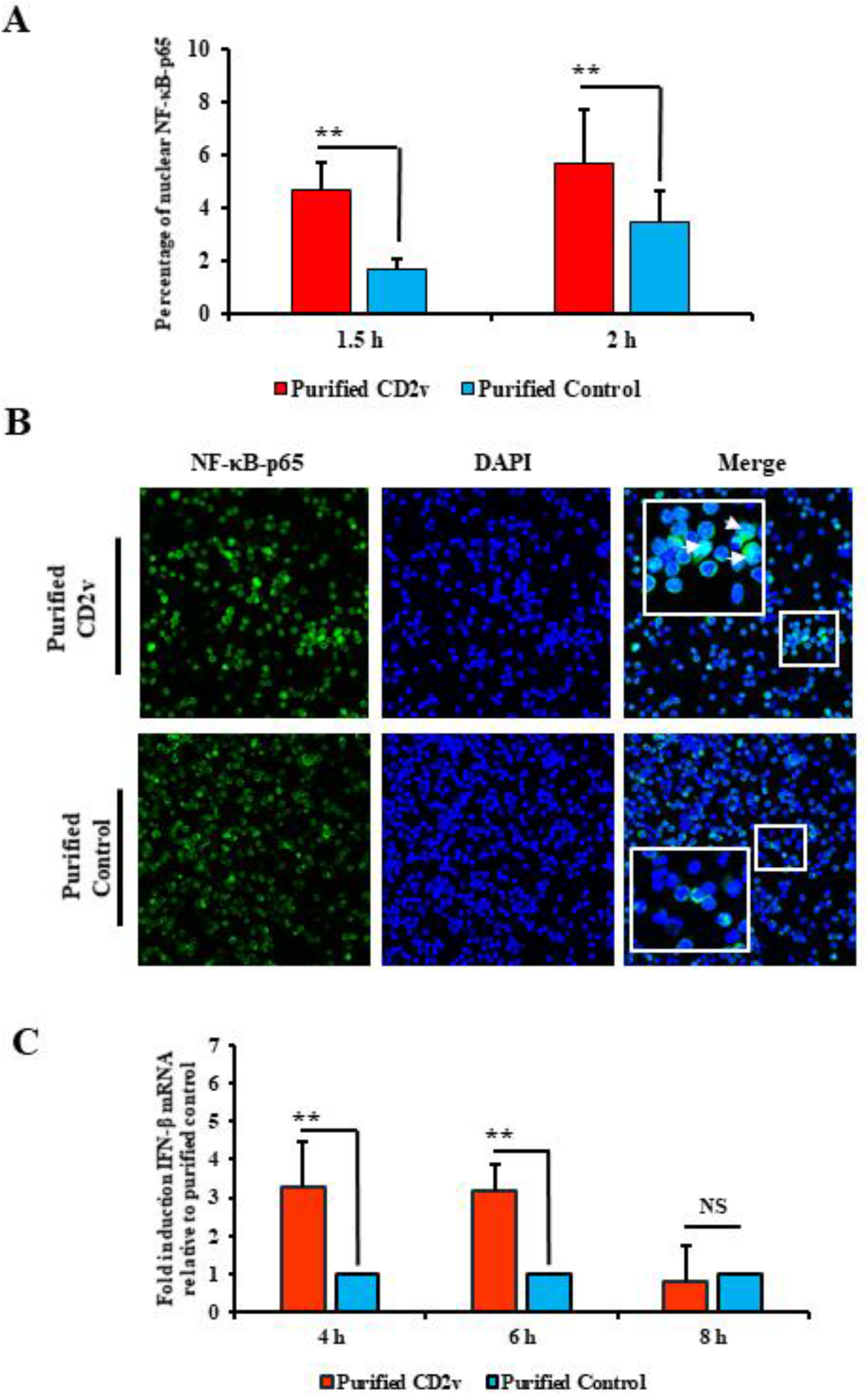
Purified CD2v induces NF-kB-p65 nuclear translocation and IFN-β transcription in swine PBMCs. Purified CD2v, purified control, and swine PBMCs were obtained as indicated in materials and methods. (A) Swine PBMCs were treated with purified CD2v or purified control, fixed at 1.5 h and 2 h post treatment, sequentially incubated with anti-NF-κB-p65 primary antibody and Alexa fluor 488-labeled secondary antibody, stained with DAPI, cytospined, and examined by confocal microscopy. Cells were counted from 15 random fields/slide (approximately 2000 cells/slide) and results are shown as percentage of cells with nuclear NF-κB-p65. Results are mean of four independent experiments (1.5 h, *P* = 0.01; 2 h, *P =* 0.012). (B) Representative confocal images of NF-κB-p65 nuclear translocation under conditions outlined in A. Green, NF-κB-p65; Blue, DAPI. Insets show magnified areas of the field. Arrows indicate nuclear NF-κB-p65. (C) IFN-β induction. Total RNA was harvested at 4 h, 6 h and 8 h post treatment and IFN-β transcription was assessed by RT-PCR. Fold changes are relative to purified control and data are mean mRNA levels of seven independent experiments (4 h, *P* = 0.016; 6 h, *P* = 0.002). (*, *P*<0.05 and **, *P*<0.01).

To investigate whether IFN-β transcription in lymphocytes is affected by CD2v, swine PBMCs were treated as above and assessed for IFN-β transcription by RT-PCR at various times post-treatment. We found that IFN-β transcription was significantly induced in CD2v-treated PBMCs at 4 h (3.3-fold) and 6 h (3.2-fold) post treatment compared to purified control (Fig. 10C).

### Supernatants from CD2v-treated swine PBMCs exhibit antiviral activity

To examine the functional significance of CD2v-induced IFN-β expression in swine lymphocytes, swine PBMCs were treated with purified CD2v or purified control for 24 h, and the supernatants collected, diluted, and used to treat fresh PK15 cells. Twenty-four hours after treatment the cells were infected with VSV^GFP^ and examined for virus replication at 16 h post infection by IFA. Supernatant collected from PK15 cell culture transfected with Poly I:C was used as positive control. Significant inhibition of VSV^GFP^ replication was observed in PK15 cells treated with supernatant from PBMCs. We observed 32.1% inhibition with undiluted supernatant, 24.4% inhibition with 1:2 dilution and 28.1% inhibition with 1:4 dilution (Fig. 11A and B).

**FIG 11.**
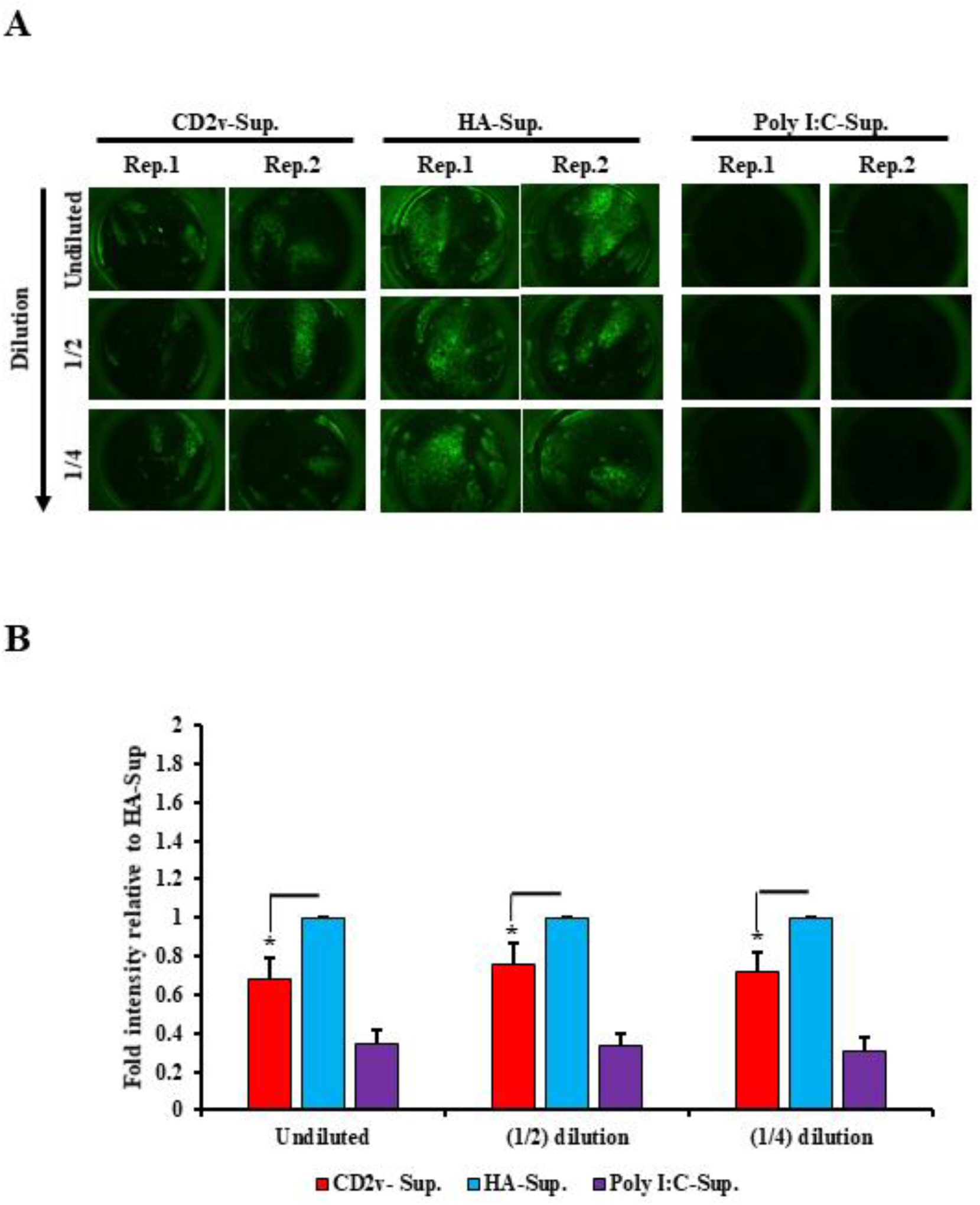
Antiviral activity of supernatants from CD2v-treated swine PBMCs. Swine PBMCs were treated with purified CD2v protein or purified control, and supernatants collected 24 h post treatment. Supernatant collected from PK15 transfected with Poly I:C (positive control). PK15 grown in 96 well plates were treated with different dilutions of PBMC supernatants for 24 h and subsequently infected with VSV^GFP^ (50 PFU/well) for 16 h. (A) Representative fluorescence images of PK15 fixed at 16 h post VSV^GFP^ infection. Green, VSV^GFP^. Rep., denotes technical replicates. (B) Intensity (mean gray value) measured by ImageJ. Fold changes are relative to purified control and data are mean of four independent experiments. *P*-values relative to purified control for undiluted, 1:2 diluted and 1:4 diluted supernatants were 0.014, 0.033 and 0.017, respectively (*, *P*<0.05).

### CD2v induces apoptosis in swine PBMCs

ASF is characterized by severe destruction of lymphoid tissue and massive lymphocyte depletion due to apoptosis (3, 7, 8, 16, 19, 20). An explanation for this critical pathogenic event is lacking. ASFV replicates in cells of the monocyte lineage, most notably macrophages, but not in lymphocytes, thus apoptosis in bystander lymphocytes is most likely due to proteins or factors secreted by infected macrophages. Based on our data, we hypothesize that CD2v released by infected macrophages induces IFN expression in bystander lymphocytes leading to apoptosis.

To investigate the effect of CD2v on PBMC apoptosis, swine PBMC cultures were treated with purified CD2v, purified control, or staurosporine (positive control) as described in the materials and methods, and caspase-3 activation and PARP1 cleavage assessed by Western blot at various times post treatment. As shown in Fig. 12 A-C, treatment of swine PBMCs with purified CD2v led to significant induction of caspase-3 activation at 18 h post treatment (1.9-fold) and PARP1 cleavage at 18 h (1.9-fold) and 24 h (1.6-fold) post treatment compared to purified control. This result indicates that CD2v treatment induces apoptosis in swine PBMC.

**FIG 12.**
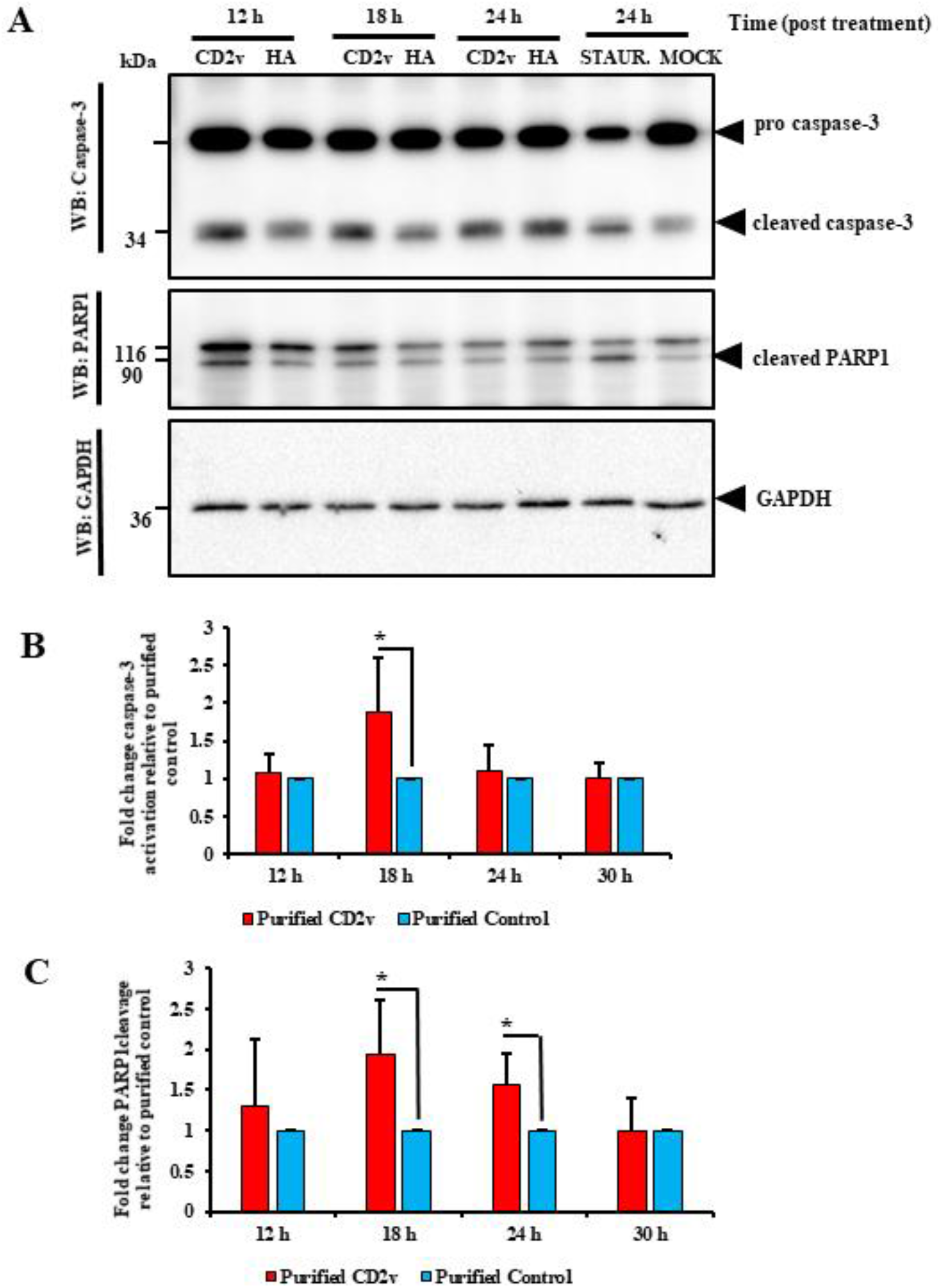
Purified CD2v induces apoptosis in swine PBMC. (A) Swine PBMC were treated with purified CD2v, purified control, or staurosporine (positive control) and whole cell lysates were obtained at 12 h, 18 h, 24 h and 30 h post treatment, resolved by SDS-PAGE, blotted, and probed with antibodies against Caspase-3, PARP1 and GAPDH. (B) Densitometric analysis showing the fold change in caspase-3 activation relative to purified control treatment, the caspase-3 results are mean values of six independent experiments (18 h, *P* = 0.042). (C) Densitometric analysis showing fold changes in PARP1 cleavage relative to purified control treatment. Results are mean values of six independent experiments (18 h, *P* = 0.026; 24 h, *P* = 0.018). (*, *P*<0.05).

### Treatment with monoclonal antibodies against ASFV CD2v inhibits CD2v-induced NF-κB activation and IFN-β transcription in swine PBMCs

Monoclonal antibodies against ASFV CD2v were generated and screened as described in materials and methods. Four anti-CD2v antibodies (A4, C4, C3 and F2) were used as a mixture to examine their reactivity against CD2v. The antibodies mix detected the full length 100 kDa CD2v species in Western blot (Fig. 13A). To confirm reactivity of antibodies, lysates from 293T cells transfected with pCD2v-HA were incubated overnight with the anti-CD2v monoclonal antibodies. Immunoprecipitation products were assessed by Western blot using anti-HA antibodies. Strong pull down of CD2v by anti-CD2v monoclonal antibodies was observed in CD2v-but not in control-transfected cells (Fig. 13B). The results show that the anti-CD2v monoclonal antibodies react with CD2v.

**FIG 13.**
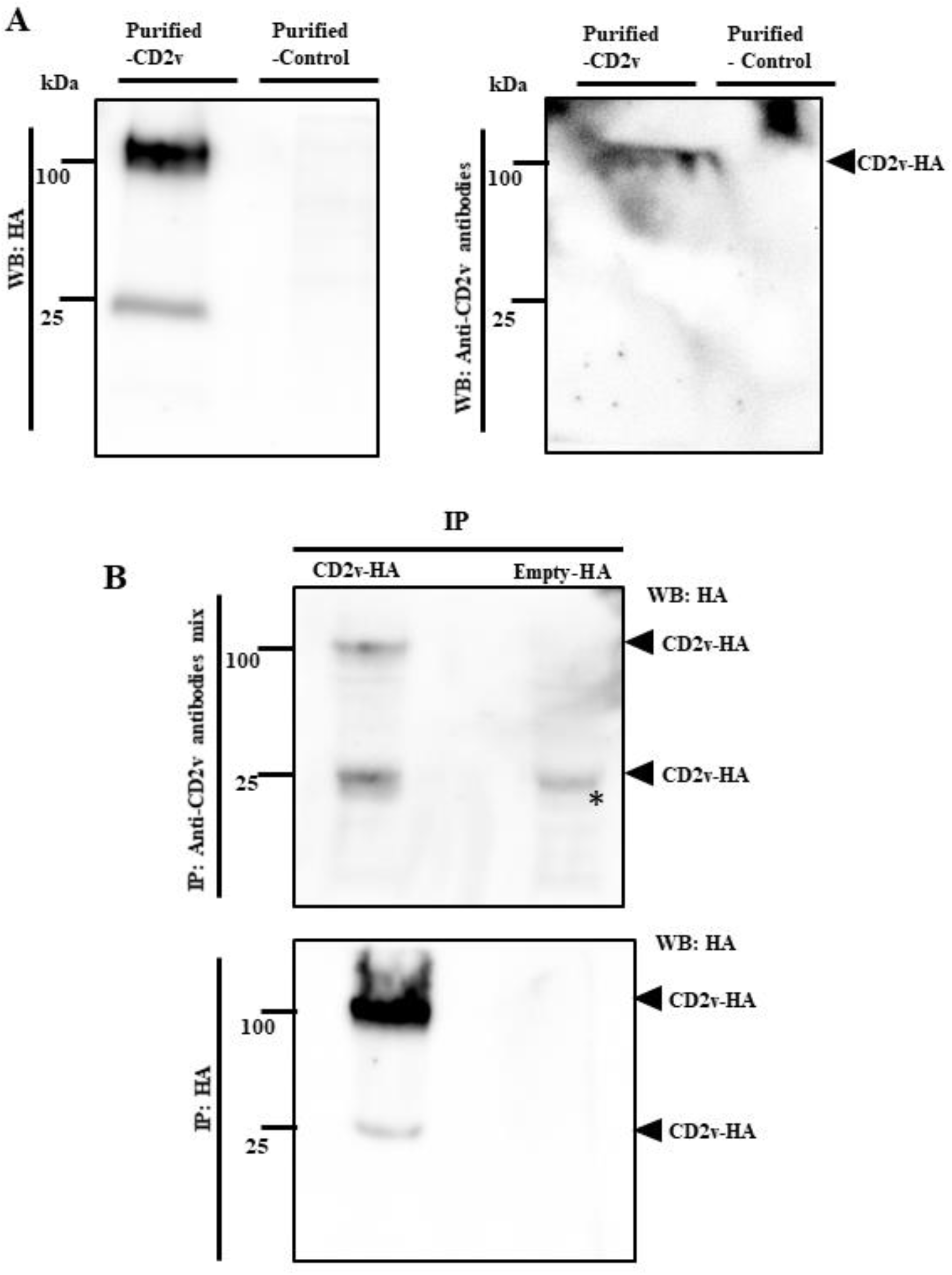
Monoclonal antibodies generated against ASFV CD2v identifies CD2v in Western blots. (A) Purified CD2v or purified control, were resolved by SDS-PAGE, blotted, and probed with anti-CD2v monoclonal antibodies (A4, C4, C3 and F2; top right) or probed with anti-HA antibody (control; top left). Results are representative of three independent experiments. (B) Whole cell extracts were immunoprecipitated with anti-CD2v monoclonal antibodies (upper blot) or anti-HA antibody (control; lower blot), resolved with SDS-PAGE, and probed with anti-HA antibody. Results are representative of three independent experiments. * denotes light chain band.

To investigate the effect of the anti-CD2v antibodies on CD2v-induced NF-κB activation in swine PBMC, purified CD2v or purified control were incubated overnight with the monoclonal antibody mix, anti-ORFV086 monoclonal antibody, or anti-IgG mouse isotype antibody control. Swine PBMCs were then treated with purified CD2v or purified control pre-incubated with the various antibodies, processed with NF-κB nuclear translocation assay as above, and analyzed by confocal microscopy. As a control for the experiment, purified CD2v or purified control without pre-incubation with antibodies was used to treat swine PBMCs. Significant inhibition of NF-κB-p65 nuclear translocation (approximately 50% reduction) was observed in PBMCs treated with CD2v previously incubated with anti-CD2v monoclonal antibodies compared to controls (Fig. 14A and B). This result supports a role of secreted CD2v in induction of NF-κB-p65 nuclear translocation in swine PBMCs.

**FIG 14.**
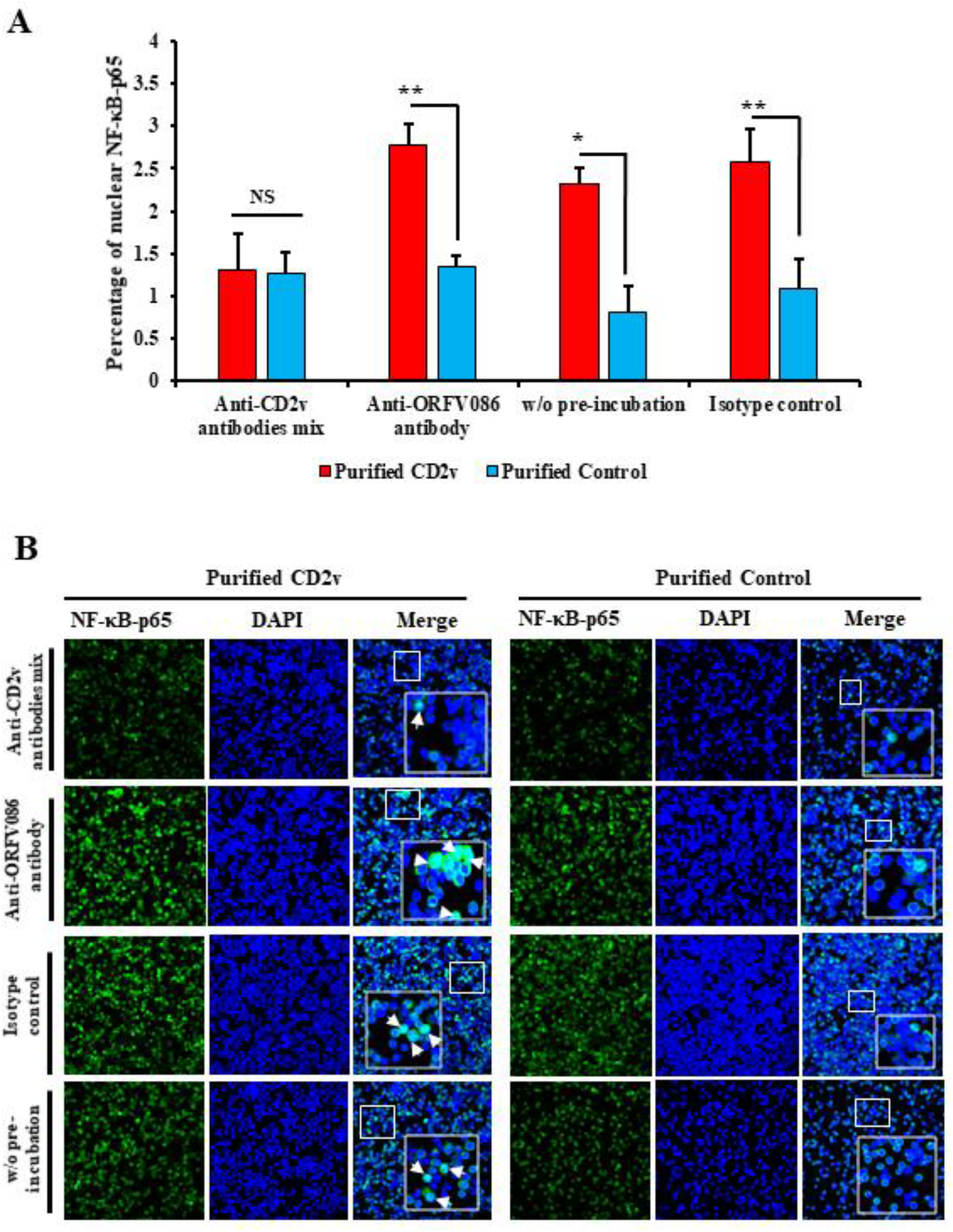
Monoclonal antibodies against ASFV CD2v inhibit CD2v-induced NF-κB activation in swine PBMCs. (A) Purified CD2v or purified control pre-incubated with anti-CD2v monoclonal antibody mix, anti-ORFV086 antibody or isotype control antibody were used to treat swine PBMCs for 1.5 h, and cells were processed for NF-κB-p65 staining by indirect immunofluorescence as explained in materials and methods. Control cells were treated with CD2v or purified control that have not been pre-incubated with antibodies. Cells were counted from 15 random fields/slide (approximately 2500 cells/slide). Results, shown as percentage of cells with nuclear NF-κB-p65, are mean values from three independent experiments (for % nuclear NF-κB-p65 for anti-CD2v antibody mix relative to anti-ORFV086, *P* = 0.012; without pre-incubation, *P* = 0.031 and for isotype control, *P* = 0.03. (*, *P*<0.05; **, *P*<0.01) (B) Representative confocal images of NF-κB-p65 nuclear translocation. Green, NF-κB-p65; Blue, DAPI. Arrows indicate nuclear NF-κB-p65. Insets show magnified areas of the field.

The effect of the anti-CD2v monoclonal antibodies on CD2v-induced IFN-β expression in swine PBMC, was investigated by pre-incubating purified CD2v or purified control with the monoclonal antibodies as above, followed by treatment of PBMC. As a control for the experiment, purified CD2v or purified control without pre-incubation with antibodies was used to treat swine PBMC. Total RNA was extracted 6 h post treatment and IFN-β transcription was assessed by RT-PCR. Significant inhibition in IFN-β transcription was observed when swine PBMCs were treated with purified CD2v pre-incubated with anti-CD2v antibody mix (1-fold) as compared to purified CD2v pre-incubated with anti-ORFV086 monoclonal antibody (1.5-fold) or without pre-incubation (1.8-fold). These results confirm a role of CD2v in the induction of IFN-β transcription in swine PBMCs (Fig. 15).

**Fig 15.**
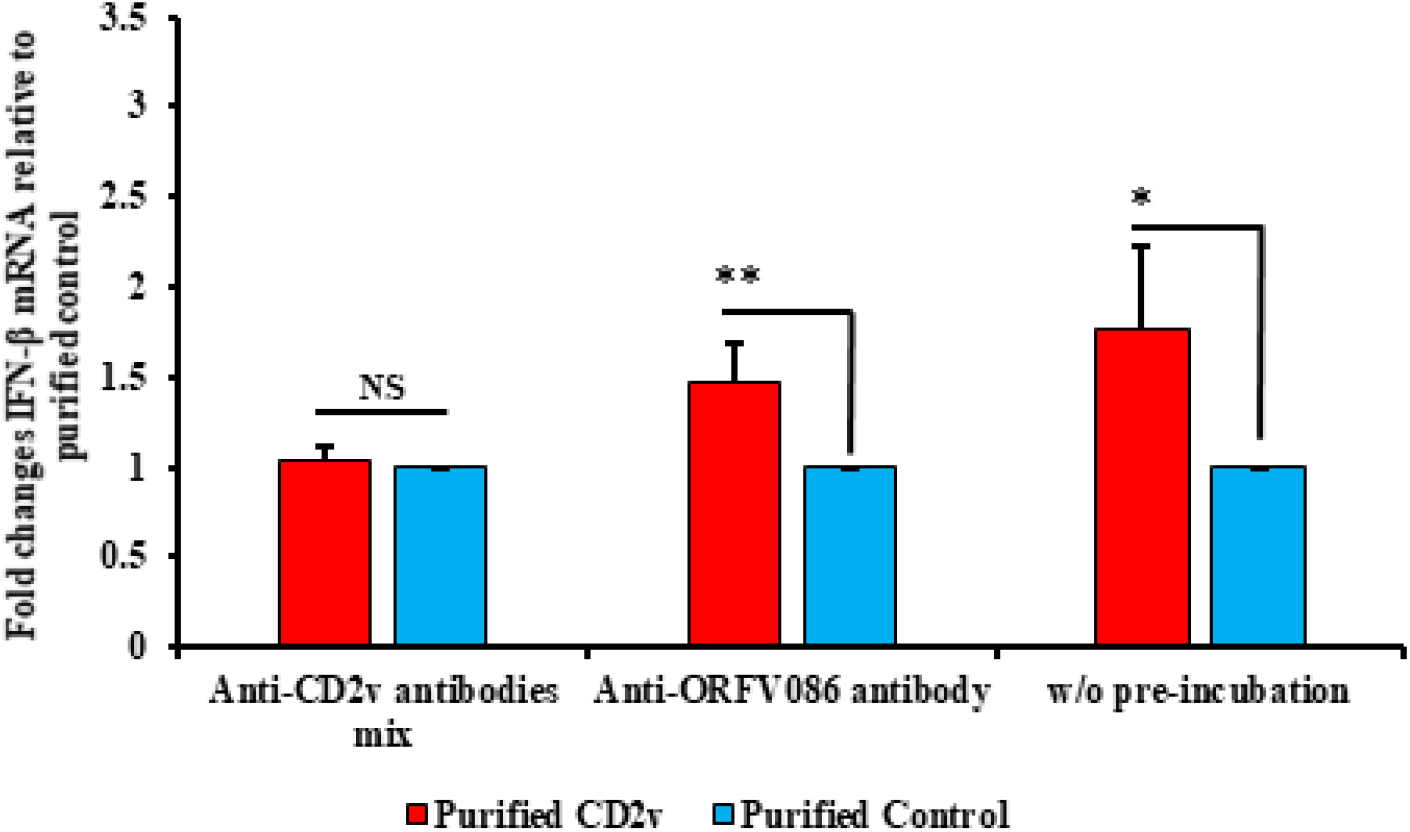
Monoclonal antibodies against ASFV CD2v inhibits CD2v-induced IFN-β transcription in swine PBMC cultures. Swine PBMCs were treated with purified CD2v or purified control pre-incubated with anti-CD2v monoclonal antibody mix or anti-ORFV086 antibody, or with the proteins without previous incubation with antibodies. Total RNA was harvested at 6 h post-treatment, and IFN-β transcription was assessed by RT-PCR. Fold changes are relative to purified control and data are mean mRNA levels from five independent experiments. *P*-value for IFN-β fold induction for anti-CD2v antibody mix compared to anti-ORFV086 is 0.014 and without pre-incubation is 0.028 (*, *P*<0.05 and **, *P*<0.01).

## DISCUSSION

A landmark of acute ASF is the severe lymphoid tissue destruction and massive lymphocyte depletion in infected pigs, which occurs as a result of bystander lymphocyte apoptosis (3, 8, 19). Since lymphocytes do not support ASFV replication, factors secreted by infected macrophages have been implicated in triggering lymphocyte apoptosis (16, 19–22). ASFV CD2v is a glycoprotein with homology to host CD2, an adhesion molecule expressed by T and NK cells (56, 57, 72). CD2v has been shown to be involved in host immunomodulation, virulence and induction of protective immune responses (55, 75, 88). Here, we show that expression of ASFV CD2v in swine cells induces NF-κB-mediated IFN expression through interaction with CD58 (Fig. 3, Fig. 5 and Fig. 9). We observed that treatment of swine PBMC with purified CD2v leads to significant IFN-β induction, NF-κB-p65 nuclear translocation (Fig. 10A and B), and caspase-3 and PARP1 cleavage, thus providing a mechanism for CD2v-induced lymphocyte apoptosis (Fig. 12).

The relationship between ASFV infection and the IFN system is complex. ASFV has evolved multiple strategies to counteract activation of the IFN and NF-κB signaling pathways (42–45). *In vitro* infection of porcine macrophages with low virulence ASFV strains induced enhanced and sustained IFN induction compared to virulent strains (46–51). However, acute ASFV infection of pigs is characterized by high level of systemic IFN production (52, 53). Viral factors involved in IFN induction and sources of IFN during acute ASF infection remain to be identified. Here, we describe an immunomodulatory function of ASFV CD2v that, by targeting lymphocytes for apoptosis, may contribute to the massive lymphocyte depletion observed *in vivo*.

CD58/LFA-3 is the natural ligand for host CD2 protein, and CD2-CD58 interaction initiates cellular kinase signaling (58–67). We show that swine CD58 interacts with CD2v, and that the interaction is involved in CD2v-mediated IFN-β induction through NF-κB-p65 nuclear translocation (Fig. 7 to 9). Since viral glycoproteins can also induce IFN-β through TLR-4/TLR-2 (76–79), involvement of these receptors cannot be ruled out. It will be interesting to determine whether the CD2v cytoplasmic domain, known to interact with the trans-Golgi network AP-1 factor and actin-binding adaptor protein SH3P7 (73, 80), plays a role in CD2v immunomodulation. In ASFV-infected macrophages, CD2v is processed and thought to be secreted (72–74). Here, we show for the first time that CD2v is released into the supernatant by CD2v-expressing cells (Fig. 2B). Previous studies have described the involvement of soluble factor/s in the inhibition of lymphocyte proliferation in PBMCs infected with ASFV or incubated with cell extracts/supernatants free of virus (54, 55) and have defined an immunomodulatory role of CD2v in inhibition of mitogen-induced proliferation of bystander lymphocytes (55). Based on our findings, secreted CD2v could be mimicking host CD2 by interacting with CD58, thus leading to induction of IFN-β and inhibition of lymphocyte proliferation. Our observation that anti-CD2v monoclonal antibodies inhibit CD2v-dependent NF-κB-p65 nuclear translocation and IFN-β induction further strengthen the immunomodulatory role of secreted CD2v (Fig. 14).

The role of CD2v in ASFV virulence is not clearly understood. Infection of pigs with a CD2v deletion mutant virus resulted in different infection phenotypes depending on the parental virus strain. CD2v deletion in the Spanish strain BA71 resulted in virus attenuation (75), and the *CD2v* gene has been found interrupted in some attenuated ASFV strains (82) suggesting a role of CD2v in virus virulence. In contrast, deletion of *CD2v* in virulent strains Malawi and Georgia 2007 did not significantly affect virus virulence in pigs (55, 81). Although not essential for viral replication in pigs, *CD2v* is critical for virus replication in the tick (83, 84). Results here support a role of ASFV CD2v in viral pathogenesis and virulence by affecting lymphocyte survival.

CD2v protein has been implicated in protective immunity. CD2v expression is required for partial protection conferred by specific vaccine constructs, and two predicted CD2v T-cell epitopes are speculated to affect protective immunity (85–87). Using ASFV inter-serotypic chimeric viruses and vaccination/challenge experiments in pigs, serotype-specific CD2v and/or C-type lectin proteins were shown to be important for protection against homologous ASFV infection (88). Others, however, have found that inoculation of pigs with the BA71 virus lacking CD2v conferred protection against parental BA71 (75). This variability likely reflects the differences in the virulence of the challenge strains used in the experiments. In conclusion, CD2v represents an important factor contributing to lymphocyte depletion observed during ASFV infection in animals (Fig. 16).

**FIG 16.**
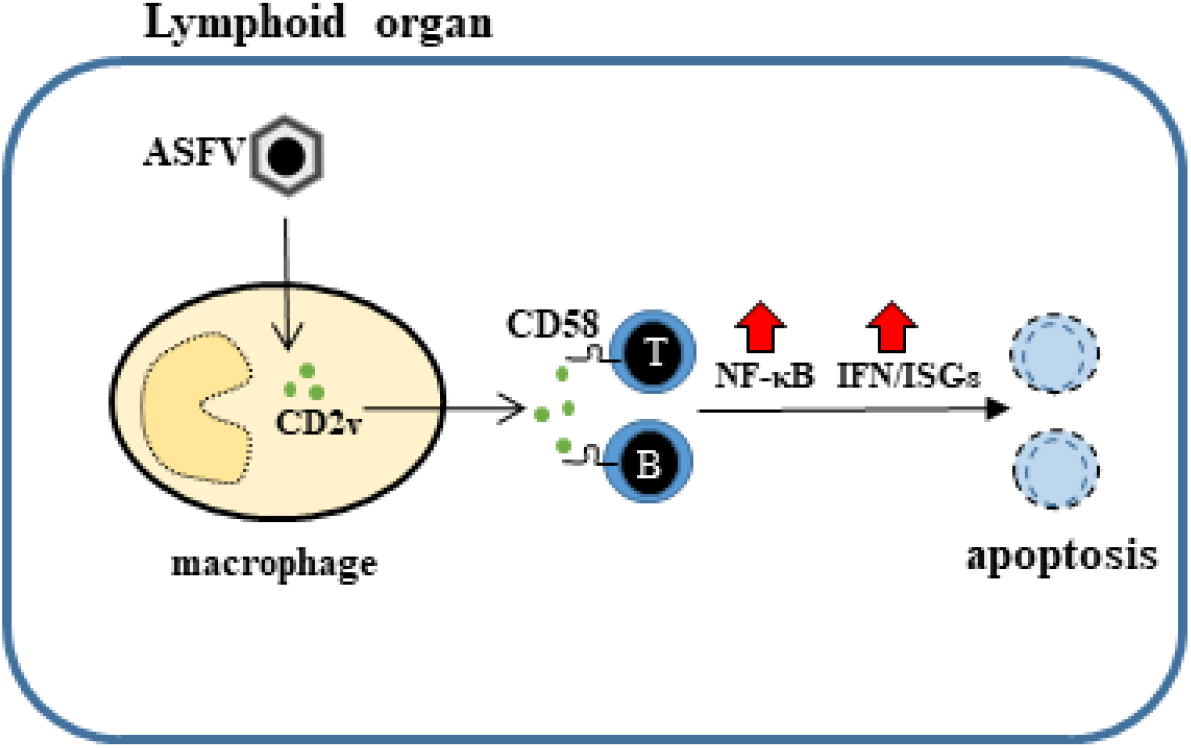
Proposed mechanism of action of ASFV CD2v. CD2v secreted from ASFV-infected macrophages interacts with by-stander lymphocytes via CD58. This interaction promotes NF-κB activation and induction of IFN-β and ISGs, which lead to lymphocyte apoptosis.

## MATERIALS AND METHODS

### Cells

Porcine kidney cells (PK15) and monkey kidney cells (Vero) were obtained from American Type Culture Collection (ATCC) and were maintained at 37° C with 5% CO2 in minimal essential medium (MEM) supplemented with 10% fetal bovine serum (FBS) (Atlanta Biologicals, Flowery Branch, GA), 2mM L-glutamine, gentamicin (50 μg/ml), penicillin (100 IU/ml), and streptomycin (100 μg/ml). Human embryonic kidney (HEK 293T) were cultured in Dulbecco’s modified essential medium (DMEM) supplemented as above.

### Plasmids and transfection

ASFV Congo strain CD2v was synthesized, cloned into pUC57 (Genscript, Piscataway, NJ) and amplified with primers CD2v-Fw (*EcoRI*) (5’-TAAGGCCTCT<u>GAATTC</u>GCCACCATGATAATTAACTTATTTTTTTAATATG-3’) and CD2v-Rv (*KpnI*) (5’-CAGAATTCGC<u>GGTACC</u>AATAATTCTATCT ACATGAATAAGCG-3’). The amplified full length *CD2v* gene was cloned into *EcoRI* and *KpnI* sites of pCMV-HA vector (Clontech, Mountain View, CA) to produce pCD2v-HA, which express C-terminally HA-tagged CD2v protein (CD2v-HA). To enhance translation efficiency, a Kozak sequence (GCCACC) was placed in front of the *CD2v* gene. The construct was sequenced to confirm integrity and fidelity.

To construct expression plasmids pORFV120-Flag and pORFV113-Flag, ORFV120 and ORFV113 coding sequences were PCR-amplified from orf virus strain OV-IA82 genome and cloned into p3xFlag-CMV-10 vector (pFlag) (Clontech).

For transfection of cells, Lipofectamine 2000 (Invitrogen) and plasmid DNA were separately diluted in Opti-MEM medium (Gibco) and incubated for 5 min. Diluted DNA was added to diluted lipofectamine 2000 (1:1ratio) and incubated for 20 min. Finally, DNA-lipid complex was added to cells and 5 h after incubation, Opti-MEM medium was replaced with 10% complete growth media.

### Hemadsorption assay

PK15 cells grown in 6-well plates were mock transfected or transfected with plasmid (p) CD2v-HA. At 24 h post-transfection (pt), culture media was removed and rinsed two times with PBS. Transfected cultures were incubated with PBS-washed 1% swine RBC overnight and observed with microscope (X100).

### Swine PBMCs isolation, freezing and culture

Swine PBMCs were obtained from swine whole blood through density gradient centrifugation using Sepmate15 (Stem cell technologies) and lymphocyte separation media (Corning), and frozen in freezing media (50% RPMI-1640, 40% FBS, 10% DMSO) as described elsewhere (68, 69). Swine PBMC were maintained at 37° C with 5% CO2 in RPMI 1640 medium (Corning) supplemented with 10% fetal bovine serum (FBS) (Atlanta Biologicals, Flowery Branch, GA), 2mM L-glutamine, gentamicin (50 μg/ml), penicillin (100 IU/ml), streptomycin (100 μg/ml) and sodium pyruvate (1 mM).

### CD2v purification

For CD2v protein purification, 293T cells were transfected with pCD2v-HA or pEmpty-HA control vector for 30 h. Cell lysates were harvested using mammalian protein extraction reagent (MPER) (Thermo Scientific) and incubated with anti-HA resin overnight using spin columns (Thermo Scientific). Next day, CD2v was eluted using HA peptide (1 mg/ml) (Thermo Scientific) in TBS buffer (Corning). Whole cell lysates obtained from two 6-well plates were incubated with 100 μl of anti-HA resin, eluted in 200 μl buffer and used for downstream process

### Monoclonal antibodies against CD2v

PK15 cells were transfected with pCD2v-HA, and whole cell lysates obtained 30 h pt. Lysates were incubated overnight with anti-HA antibody (Cell Signaling Technology) at 4° C, and immunoprecipitated using 50 μl of protein G agarose bead slurry (Millipore). Monoclonal antibodies recognizing CD2v-HA were generated by immunizing BALB/c mice with immunoprecipitated CD2v-HA (100 μg/dose). To generate hybridomas, splenocytes from immunized mice were harvested and processed according to the company’s protocol (STEMCELL technologies/ClonalCell-HY Hybridoma Cloning Kit). Clones were screened for reactivity against CD2v-HA by IFA using CD2v-HA-transfected PK15 cultures. Monoclonal antibody titers were estimated by IFA using goat anti-mouse Alexa fluor 488 or 594 as secondary antibody. Mice were maintained at the University of Nebraska-Lincoln (UNL) in accordance with the guidelines of UNL Institutional Animal Care and Use Committee (IACUC) and used in accordance with the guidelines of the committees.

### Western Blot

Fifty μg of whole protein cell extracts or 50 μl of cleared culture supernatant were resolved by SDS-PAGE, blotted to nitrocellulose or PVDF membranes and probed with primary antibody against HA (Cell Signaling Technology), caspase-3 (9662S; Cell Signaling Technology), PARP1 (sc-53643; Santa Cruz) or glyceraldehyde-3-phosphate dehydrogenase (GAPDH) (sc-25778; Santa Cruz). The blots were developed with appropriate HRP-conjugated secondary antibodies and chemiluminescent reagents (Thermo Scientific), and imaged with FluorChem R (Protein Simple). Densitometric analysis was performed using ImageJ software and all readings were normalized to GAPDH values.

### Immunoprecipitation

293T cells were transfected with pCD2v-HA, and whole cell lysates in MPER lysis buffer (Thermo Scientific) were obtained at 30 h pt and incubated overnight with the anti-CD2v monoclonal antibody mix or anti-HA antibody (3724; Cell Signaling Technology). Pull down products were obtained by eluting in Laemmli buffer (Bio-Rad) and analyzed by Western blot using anti-HA antibody (2367; Cell Signaling Technology).

### CD58 siRNA knock-down

To investigate involvement of CD58-CD2v interaction in CD2v-mediated IFN-β induction, siRNA knock down experiments were performed using CD58 sense (S) (CUUCCAGAGCCAGAACUAU) and anti-sense (AS) (AUAGUUCUGGCUCUG GAAG) siRNA duplex (Sigma Aldrich). PK15 cultures were transfected with CD58 siRNA (15 nM) and mission siRNA transfection reagent (Sigma) following the manufacturer’s protocol, and transfected 24 h later with pCD2v-HA or control pEmpty-HA for 6 h. CD58 knock-down was assessed by comparing transcript levels between cultures transfected with one MISSION siRNA Universal Negative control (Sigma aldrich) and CD58 siRNA-transfected cultures using RT-PCR. SYBR primers for swine CD58 were sCD58 FW 5’-ACTTAAACACTGGGTCGGGC-3’ and sCD58 RV 5’-AAGCTGCAAGGATCAGGCAT-3’.

### Real-time PCR

Interferon-β (IFN-β) and interferon stimulated genes (ISGs) transcription were assessed in PK15 cells transfected with pCD2v-HA or control plasmids, and in swine PBMCs treated with purified CD2v or purified control. Total RNA was harvested at various times post transfection/treatment with RNA-extraction kit (Zymo) and reverse transcribed with MLV-RT (Invitrogen) as previously described (70). IFN-β and ISGs mRNAs were quantified using ABI and QuantStudio-3 Real time PCR system (Applied Biosystems, Foster city, CA), Power SYBR Green PCR Master Mix (Applied Bio) and primers sIFNβFw (5’-AGTGCATCCTCCAAATCGC T-3’) and sIFNβRv (5’-GCTCATGGAAAGAGCTGTGGT-3’) for IFN-β mRNA, and sMX1Fw (5’-GGCGTGGGAATCAGTCATG-3’), sMX1Rv (5’-AGGAAGGTCTATGAGGGTCAGATCT-3’), sOAS1Fw (5’-GAGCTGCAGCGAGACTTCCT-3’) and sOAS1Rv (5’-TGCTTGACAAGGCGGATGA-3’) for ISGs *MX1* and *OAS*. Fold change was calculated by comparison to Empty-HA for each time points. Experiments were conducted with biological triplicates and at least three technical replicates.

### NF-κB nuclear translocation assay

Involvement of NF-κB in CD2v-mediated upregulation of IFN-β transcription was assessed by nuclear translocation assays in 1-PK15 cells transfected with pCD2v-HA or control plasmids pORFV120-Flag and pORFV113-Flag, and 2-swine PBMCs treated with purified CD2v or purified control. Cells were fixed at various times post-transfection or –treatment, fixed with 2%-4% paraformaldehyde (PFA) (15 min), and permeabilized with 0.2% triton X100 (10 min). PK15 cells were incubated overnight at 4° C with primary antibody against HA (2367S; Cell Signaling Technology), control Flag (A00187; Genscript) and total NF-κB (8242; Cell Signaling Technology), and PBMCs with antibody against total NF-κB. PK15 cells were washed with PBS and incubated with secondary antibodies, goat anti-mouse Alexa fluor 488 (A11029; Thermo Scientific) to detect CD2v, ORFV120 and ORFV113, and goat anti-rabbit Alexa fluor 594 (A11012; Thermo Scientific) to detect total NF-κB; for PBMCs, goat anti-mouse Alexa fluor 488. PBMCs were deposited on slides using Shandon cytospin 2 centrifuge (1500 rpm, 1 min). Nuclei were counterstained with DAPI. Images were obtained with a A1 Nikon confocal microscope, and the number of cells exhibiting nuclear NF-κB staining was determined in randomly selected fields and results were expressed as mean percentage of cells with nuclear NF-κB over three independent experiments.

To assess whether inhibition of NF-κB affects IFN-β induction by CD2v, PK15 cells were pretreated with the NF-κB inhibitor parthenolide (InvivoGen) at 1μM final concentration or vehicle control (DMSO) for one hour and transfected with pCD2v-HA in presence or absence of parthenolide (1 μM). At 3 h pt, cultures were fixed, permeabilized, and incubated with primary antibodies against HA and total NF-κB as above, washed, and incubated with secondary antibodies, goat anti-mouse Alexa fluor 488 for CD2v and goat anti-rabbit Alexa fluor 594 for total NF-κB. Cells were counterstained with DAPI and percentage NF-κB nuclear translocation in CD2v expressing cells in presence or absence of inhibitor were compared over three independent experiments.

The role of CD2v-CD58 interaction in induction of NF-κB nuclear translocation was examined by transfecting PK15 cells with CD58 siRNA or siRNA universal negative control for 24 h, and with pCD2v-HA 24 h later for 3 h. Cultures were fixed, permeabilized and processed for HA and total NF-κB staining as described above. Percentage of NF-κB nuclear translocation in CD2v-expressing cells in which CD58 was knocked down were determined and compared in three independent experiments.

To investigate the effect of anti-CD2v monoclonal antibody mix in CD2v-induced NF-κB activation in swine PBMCs, purified CD2v or purified control were incubated overnight at 4° C with anti-CD2v monoclonal antibodies or anti-ORFV086 antibody or anti-IgG mouse isotype antibody control. PBMCs grown in 96-well plates were treated with pre-incubated CD2v or non-preincubated CD2v control, processed as above for NF-κB nuclear translocation 1.5 h post treatment, cytospinned, and examined under the confocal microscope. Fields were randomly selected and scored for mean percentage of cells containing nuclear NF-κB over three independent experiments.

### Interferon Bioassay

To investigate the functional significance of CD2v-induced IFN-β and ISGs expression, the antiviral state of cells was examined using an IFN bioassay. PK15 cells were transfected with pCD2v-HA or control plasmids pEmpty-HA or pORFV120-Flag, infected at 12 h, 24 h and 30 h pt with reporter vesicular stomatitis virus expressing GFP (VSV^GFP^; 50 PFU/well), fixed with 4% PFA at 16 h post infection, and examined by IFA in a Leica DMI 4000B.

Supernatant obtained from PK15 cultures transfected with pCD2v-HA or control plasmids pEmpty-HA or pORFV120-Flag or Poly I:C (+ control) were serially diluted and used to treat PK15 cells. At 30 h post-treatment, cultures were infected with VSV^GFP^ (50 PFU/well), and virus replication was assessed 16 h post-infection.

The antiviral state was assessed in swine lymphocytes as follows. Supernatants from swine PBMCs treated with purified CD2v or purified control for 24 h or supernatant obtained PK15 culture transfected with Poly I:C (+ control) were serially diluted and used to treat fresh PK15 cells. PK15 cells were infected with VSV^GFP^ (50 PFU/well) 24 h post treatment, and virus replication was examined 16 h post infection as described.

### Flow cytometry

PK15 cells were transfected with pCD2v-HA or control plasmids, and then infected with VSV^GFP^ at 12 h, 24 h and 30 h pt. At 16 h post infection, cells were trypsinized, fixed with 2% PFA, and washed. GFP mean fluorescence intensity (MFI) was measured using Cytek Aurora flow cytometer (Cytek Biosciences, Fremont CA).

CD2v activation of the NF-κB pathway was further examined using flow cytometry assay. PK15 cells were transfected with pCD2v-HA or pORFV113-Flag, fixed with 2% PFA at 3 h pt, permeabilized with 0.2% Triton-X100 and incubated with anti-HA (2367S; Cell Signaling Technology), anti-Flag (A00187; Genscript) or anti-phosphorylated NF-κB (S536) (3033S; Cell Signaling Technology) for 45 min on ice. Secondary antibodies were goat anti-rabbit Alexa fluor 488 (A11008; Thermo Scientific) for pNF-kB (S536) and goat anti-mouse Alexa fluor 647 (A21236; Thermo Scientific) for CD2v and ORFV113. pNF-κB (S536) mean fluorescence intensity (MFI) was examined in cells expressing CD2v or ORFV113.

### Co-immunoprecipitation

To study the interaction between CD2v and CD58, PK15 cells were co-transfected with pEmpty-HA and pCD58-Flag or pCD2v-HA and pCD58-Flag. Whole cell extracts were prepared at 8 h pt using RIPA lysis buffer (Thermo Scientific). Reciprocal co-immunoprecipitations were performed using active motif Co-IP kit (Active Motif, Carlsbad, CA) in moderate stringency buffer (IP High buffer [Active Motif] and Protease inhibitor cocktail [Sigma], without detergent and salt) following the manufacturer’s protocol. Whole cell extracts were incubated with HA (3724S; Cell Signaling Technology) and Flag (A00187; Genscript) antibodies overnight at 4° C, and then pre-washed with 50 μl of protein G agarose bead slurry (Millipore) at 4° C for 2 h. Beads were washed four times with moderate stringency buffer and bound proteins were eluted in Laemmli buffer. Whole cell protein extracts and immunoprecipitated products were examined by SDS-PAGE-Western blot with the appropriate antibodies.

Interaction of CD2v with endogenous human CD58 was evaluated in 293T cells transfected with pCD2v-HA. Reciprocal co-immunoprecipitation was performed using anti-HA (3724S; Cell Signaling Technology) and hu-CD58 (sc20009; Santa Cruz) antibodies following the manufacturer’s protocol (Active motif Co-IP kit). Immunoprecipitated products were examined by SDS-PAGE-Western blot with the appropriate antibodies (7074; Cell Signaling Technology, 7076; Cell Signaling Technology).

### Co-localization assays

CD2v-CD58 interaction was examined by confocal microscopy. PK15 cells were co-transfected with pCD2v-HA and pCD58-Flag, fixed with 4% PFA 24 h pt, permeabilized with 0.2% Tritron-X 100 and treated with anti-HA (3724S; Cell Signaling Technology) or anti-Flag (A00187; Genscript) primary antibodies. Cells were then washed with PBS and incubated with goat anti rabbit Alexa fluor 594 (A11012; Cell signaling Technology) and anti-mouse Alexa fluor 488 goat (A11029; Cell Signaling Technology) secondary antibodies for 1 h. Nuclei were stained with DAPI, and images obtained using A1 Nikon confocal microscope. Similarly, co-localization of CD2v with endogenous human CD58 was investigated in 293T cells. Cells were transfected with pCD2v-HA and IFA was performed using anti-HA (3724S; Cell Signaling Technology) and anti-CD58 (sc20009; Santa Cruz) primary antibodies, and goat anti rabbit Alexa fluor 594 (A11012; Cell Signaling Technology) and anti-mouse Alexa fluor 488 goat (A11029) secondary antibodies.

### Statistics

All statistical analyses were performed using student’s t test. Statistically significant difference were indicated as *, *P*<0.05; **, *P*<0.01 and NS, not significant.

## ACKNOWLEDGEMENTS

This work was funded by National Pork Board (grant 13-102) and by the USDA National Institute of Food and Agriculture (grant 2013 – 67015-21335).

We thank Hiep Vu (Nebraska Center for Virology, University of Nebraska-Lincoln) for his advice and assistance in generating monoclonal antibodies, Fernando A. Osorio (Nebraska Center for Virology, University of Nebraska-Lincoln) for laboratory support, James F. Lowe (College of Veterinary medicine at University of Illinois at Urbana-Champaign) and Anna Carol Dilger (Department of Animal Sciences at University of Illinois at Urbana-Champaign) for providing swine blood.

---

**FIG S1.**
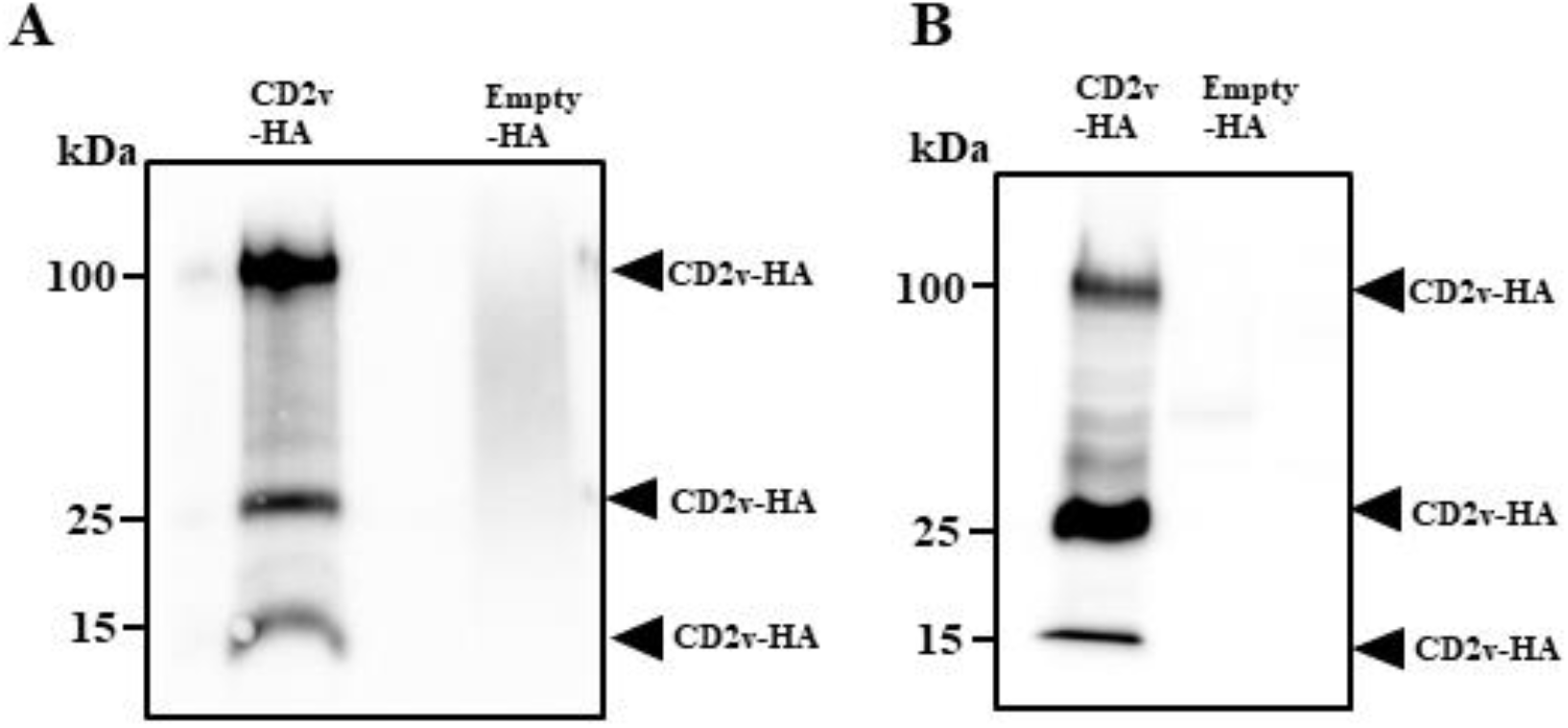
Expression of CD2v in 293T and Vero cells. 293T (A) and Vero cells (B) mock transfected or transfected with plasmid pCD2v-HA were harvested at 24 h post transfection. Total cell protein extracts were resolved by SDS-PAGE, blotted and incubated with antibodies against HA. Results are representative of two independent experiments.

**FIG S2.**
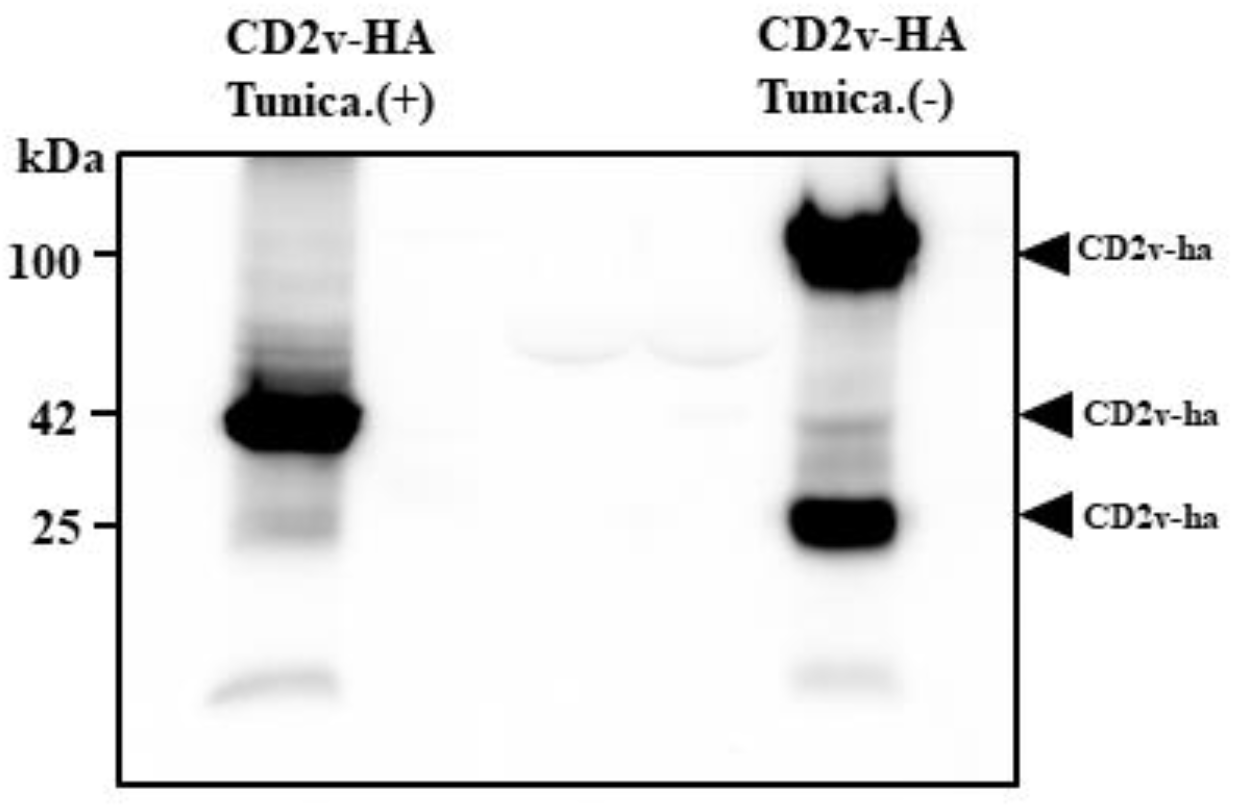
CD2v expression in presence of tunicamycin. Pk15 cells transfected with pCD2v-HA. Five hours post-transfection, media was replaced with complete growth media containing 1μg/ml tunicamycin and incubated for 24 hr. Total cell extracts were resolved by SDS-PAGE, blotted and incubated with antibodies against HA. Results are representative of two independent experiments.

